# Molecular-level modulation of the SAM-I riboswitch drives global dynamical shifts

**DOI:** 10.64898/2026.03.23.713157

**Authors:** Vrinda Nair, Mohammadreza Niknam Hamidabad, Deniz Meneksedag Erol, R.A. Mansbach

## Abstract

There has been a surge in antibiotic resistance in recent years, making traditional protein-targeting antibiotics less effective against key pathogens. RNA has recently emerged as a potential target for antibiotics due to its involvement in crucial microbial functions. It is possible to expand the range of therapeutic targets by using RNA-based therapies, but it remains necessary to improve the molecular-level understanding of interactions between RNA and known and potential binders. The SAM-I riboswitch, which controls the transcriptional termination of gene expression involved in sulfur metabolism in most bacteria, is an excellent ligand target. Thus, understanding its behavior with and without ligand complexes would be very helpful for drug design applications. In this manuscript, we studied the interactions between the SAM-I riboswitch and its natural ligand, SAM, which controls riboswitch function, and compared those interactions to its interactions with the very similar small molecular SAH, which does not control riboswitch function, and to its interactions with a potential binder JS4, identified via virtual screening. From our simulations, we gain a deeper understanding of small molecule interactions with the SAM-I riboswitch on both a local and a distal level. The results reveal how differently the small molecules (SAM, SAH and JS4) bind to and potentially induce conformational changes in the riboswitch. Our findings offer valuable insight into the molecular mechanisms underlying riboswitch RNA-ligand interactions for the design of more effective RNA-targeting therapeutics.

## 1. INTRODUCTION

The escalating challenge of antimicrobial resistance highlights the need to identify and pursue drug targets beyond conventional proteins. In particular, non-coding RNAs are emerging as promising therapeutic targets as they play critical roles in fundamental cellular processes such as cell division and DNA damage repair^1^. For example, aminoglycosides, which inhibit protein synthesis by targeting the 16S rRNA A-site of the 30S ribosomal subunit, have demonstrated efficacy in treating the intractable infections arising from Gram-negative bacteria^2^.

Riboswitches are multi-domain units of non-coding RNA that recognize binding partners through their *aptamer domain* and cause distal conformational shifts in their *expression domain* in response. This “switching” behavior regulates gene expression, including transcription, termination, and translation^3^. More than 50 distinct classes of riboswitches have been discovered since the original identification of the adenosylcobalamin riboswitch^4^ in 2002^5,6^. Riboswitches have been found in the genome of approximately 700 bacterial species. Because of their widespread nature in bacteria, it has been suggested that riboswitches might be a potential molecular target for new types of antibiotics^5^.

In particular, the S-adenosylmethionine-I (SAM-I) riboswitch is understood to regulate the biosynthesis of its binding partner S-adenosylmethionine (SAM), which is involved in regulating a number of genes involved in important cellular processes. SAM-I riboswitches have been identified in Gram-positive bacteria such as Bacillus anthracis and Bacillus subtilis, located upstream of genes that regulate essential cellular processes such as cysteine biosynthesis, sulfur metabolism, methionine biosynthesis and the biosynthesis of SAM itself ^6,7,8^. The current understanding is that SAM-I riboswitch regulates SAM biosynthesis by modulating transcription termination. At low cellular concentrations of SAM, the riboswitch exists in a dynamic conformational ensemble in which the expression platform can often take on a structure that permits gene expression, termed an antiterminator loop. At higher concentrations, SAM binds at the junction of the P1 and P3 loops of the aptamer domain (**see Fig. 1**), inducing a conformational shift in which the P1 loop interacts with the expression platform to stabilize a different structure that limits gene expression, termed a terminator loop^9^. Experimental studies have shown that the interaction between the riboswitch and the ligand is highly specific. S-adenosylhomocysteine (SAH), which is very similar to SAM but has a neutral sulfur because it lacks a methyl group bonded to the sulfur, was reported to bind the riboswitch; however, without inducing transcription termination^10^. This and proximity in the crystal structure suggests that the specificity for SAM potentially arises from a strong electrostatic interaction between the positive sulfonium ion of SAM and the electronegative carbonyl oxygens of uracil residues U7 and U88^10^. Accordingly, a possible metric for evaluating a small molecule as a SAM mimic is whether it engages the SAM-I riboswitch in a binding mode that closely resembles that of SAM. In contrast, ligands that bind in a manner more similar to SAH may fail to induce the conformational changes required for transcription termination. To rigorously assess these distinctions, both interactions must be dynamically characterized in atomistic detail.

**Figure 1:**
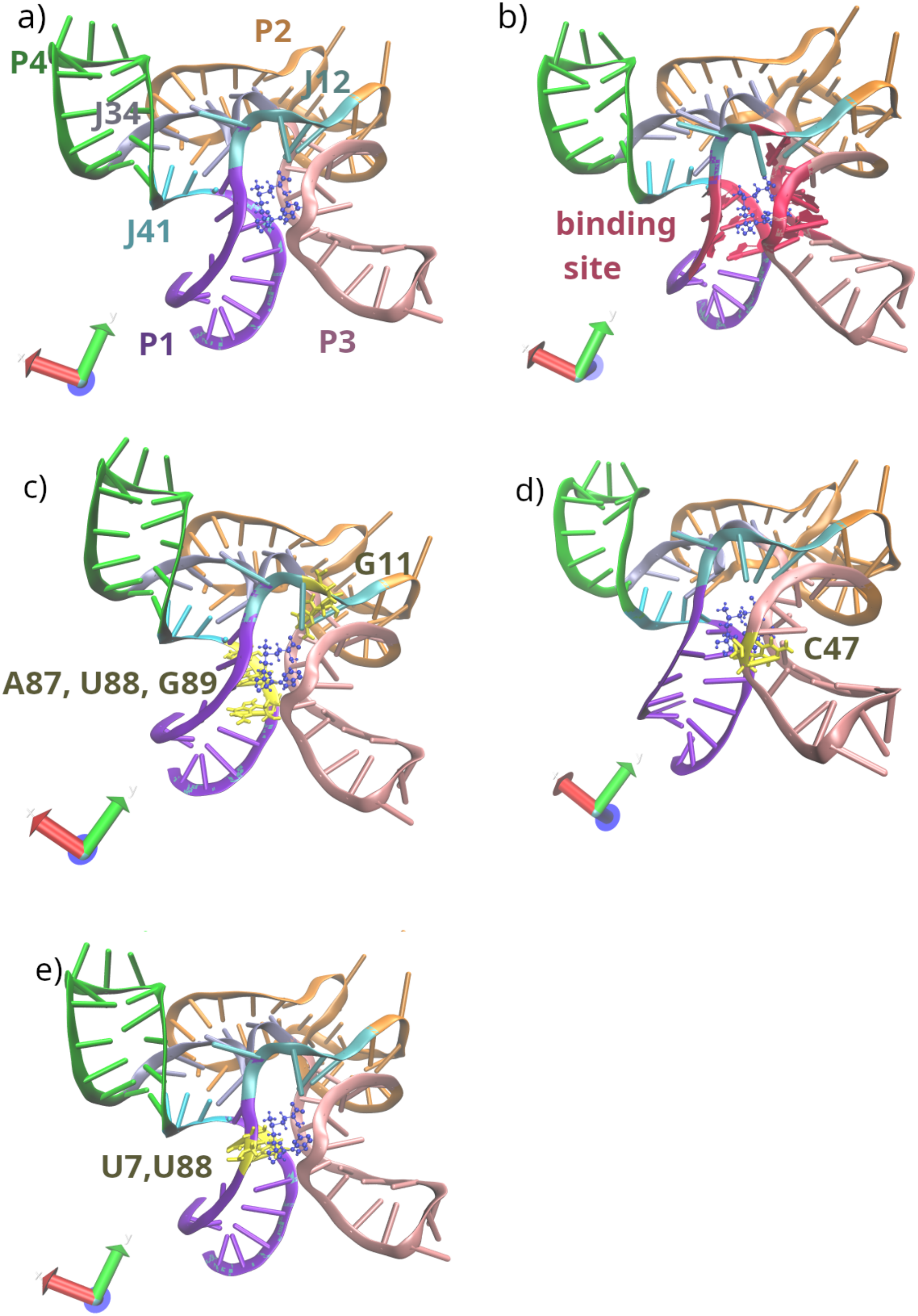
Aptamer domain of the SAM-I riboswitch with bound SAM ligand (dark blue) in the bipartite pocket formed between the minor grooves of helices P1 and P3 (a) with regions colored and labeled (b) with binding site highlighted in red, (c) with residues involved in ligand hydrogen bonding highlighted, (d) with residue C47, which is involved in pi-pi stacking interactions, highlighted, and (e) with residues U7 and U88 highlighted. The image is created from the PDB ID 3GX5 with VMD.

Molecular modeling studies of the SAM-I riboswitch are limited; correspondingly, so are direct evaluations of its dynamics at a molecular level. Elkholy et al.^7^ applied molecular docking, short MD simulations and mm-PBSA calculations to identify potential inhibitory compounds with desirable properties. Huang et al.^12^ carried out microsecond-long simulations of both aptamer and expression domains in the presence and absence of SAM, starting from structural models in an “intermediate” conformation between antiterminator and terminator. They showed that SAM binding tends to induce further formation of base pairs associated with the P1 loop of the terminator structure, thus favoring conformations associated with the termination of gene expression. Thus far, no molecular modeling study has considered SAH binding as well as SAM binding.

This work investigates the interactions of known and potential binders of the SAM-I riboswitch in atomistic detail. We simulated the aptamer platform and P1 loop of the SAM-I riboswitch in complex with the SAM ligand known to control P1 loop zipping, with the SAH ligand that is known to bind but not to control gene expression, and with a potential ligand JS4 (found from virtual screening). Through a total of 25 microseconds’ worth of molecular dynamics simulations, we assessed the conformations of the riboswitch with and without the small molecules, investigated molecular interactions between small molecules and RNA over time, and probed the contributions of the small molecules to the stability of riboswitch RNA. Through a combination of local contacts assessments and more distal network analyses, we investigated similarities and differences between the different systems. Our work sheds light on the molecular-level characteristics of SAM-I riboswitch-ligand binding and the effects of different ligands on the local and distal shape of the RNA aptamer platform, which is important for later development of small molecules as potential SAM-I inhibitors.

## 2. METHODOLOGY

### 2.1 Overview

We first performed virtual screening to identify a potential ligand binder to SAM. After identifying the most promising candidate with molecular docking, we proceeded to perform MD simulations of the SAM-I riboswitch in the structure associated with ligand binding in complex (i) with the SAM ligand, (ii) with the SAH ligand, and (iii) with the identified top hit from virtual screening, as well as (iv) with no ligand bound. We hypothesized that when complexed with SAM, we would observe lower flexibility in the P1 loop region compared to the riboswitch with no ligand bound or when bound to SAH, since the P1 loop region is primarily stabilized in the presence of SAM as part of stabilization of the terminator loop, while it exists in an “unzipped” flexible state in the ligand’s absence.. In addition to proving or disproving the hypothesis regarding the more distal effect of SAM binding on the P1 region and characterizing them in detail with a network analysis, we characterized the local interactions of the ligand in the binding site and studied how tightly bound the different ligands were and what interactions stabilized their binding, as well as how persistent or transient the stabilizing interactions were.

### 2.2 Virtual screening via molecular docking for ligand candidate identification

As a first step, we performed virtual screening to identify a ligand with the potential to bind the SAM-I riboswitch. We curated a small dataset of potential RNA binders from two databases: (i) R-BIND (The RNA-Targeted Bioactive ligand Database)^13^ and (ii) Hariboss (Harnessing RIBO nucleic acid- small molecule structure)^14^. Both the databases (R-BIND and Hariboss) specifically focus on small molecules that are already known binders of RNA structures. We downloaded the complete dataset of small molecules from the R-BIND and Hariboss database, then we excluded all entries with molecular weights of more than 800 g/mol and considered only small molecules that had associated RNA-molecule bound structures acquired by X-ray crystallography. Our final dataset consists of 150 unique small molecules curated from R-BIND and Hariboss, which were downloaded in the SDF format. After curating our data from both databases, we cross-checked our curated dataset to find if there were any duplicates. We then docked our curated small molecules with the SAM-I riboswitch RNA (PDB ID: 3GX5). Molecular docking was performed using FITTED (Flexibility Induced Through Targeted Evolutionary Description)^15^, a suite of automated computational tools designed for docking flexible or rigid ligands into flexible or rigid target molecules. Following target preparation and binding site identification, we performed flexible docking with a ligand cut-off of 7Å for our screening. We did five runs of docking for each small molecule to our RNA target, generating five poses per ligand (See **Section S1.1** for more details.)

### 2.3 Molecular dynamics simulations

All-atom MD simulation was carried out using the GROMACS software (version 2023.2)^16^ with the AMBERFF14SB force field^17^ and the OL3 parameter set. The OL3 parameter set is a specialized force field parameter set that refines the glycosidic torsion angle χ to accurately model single-stranded RNA^18,19^. Previous work has adopted this combination of force fields for accurate modelling of stability and flexibility of RNA structures^20^. It is important to note that RNA simulations remain sensitive to force field limitations and parameter choices; we choose to use a combination that is validated for RNA and has been employed previously in the context of assessment of RNA complex conformational stability and flexibility as we are particularly focused on the flexibility of the different domains as a proxy for potential conformational changes.

We employed the TIP4PEW water model, as has been done previously^21^; it provides a balance of accuracy and computational efficiency^22,23^. To model Mg2+ ions for AMBER force fields, a 12-6-4 LJ (Lennard-Jones)-type nonbonded model was employed, the standard in CHARMM-GUI. This approach is used to overcome the known deficiencies of standard 12-6 LJ potentials in representing the strong polarization and specific coordination geometry of divalent cations^24,21^.

All studies were performed using the crystal structure of the SAM-I riboswitch RNA with PDB ID 3GX5, which is co-crystallized with SAM (ligand), Mg (magnesium) and K (potassium) ions, but the SAM ligand was removed except when studying the native ligand-bound system. This RNA structure has two point mutations at U34C (mutation in the P2 loop) and A94G (a mutation at the 3’ terminus of the sequence), which were done to improve the resolution of the RNA structure, which is a common strategy used in crystallography^10^. Despite these mutations, we employed this structure for direct comparison with the experimental study which produced it, in which the authors reported very similar binding structures for SAM and SAH but experimental isothermal calorimetry highlighted the potential role of U7 and U88 in discriminating between the ligands. In total, five systems were simulated: i) original 3GX5 with experimentally co-crystallized SAM, ii) 3GX5 with SAM docked, iii) 3GX5 with docked SAH, iv) 3GX5 with docked JS4 (potential binder), and v) 3GX5 with no ligand in the binding site. We simulated both the original structure and a structure with docked SAM to control for potential differences in initial configurations, as we only had access to docked conformations for ligands other than SAM.

#### 2.3.1 RNA-ligand complex preparation for MD simulation

We used the CHARMM-GUI web server^25,26,27^ to prepare our RNA-ligand complexes for MD simulation. Charges were set assuming pH 7, with the terminal group patching as 5TER (5’-terminal hydroxyl patch)- 3TER (3’-terminal hydroxyl patch), that is with nucleic acid structures hydroxyl groups patched to the terminal ends to reflect the naturally protonated states of the terminal hydroxyl groups. Small molecule parameterization was performed (in CHARMM-GUI) using the Antechamber module with the General Amber Force Field 2 (GAFF2) atom typing scheme^28^. Partial atomic charges for the ligand were assigned using the AM1-BCC (Austin Model 1-Bond Charge Corrections) charge method^29,28^. The RNA models were prepared for molecular dynamics simulations by first neutralizing the system with Mg²⁺ counterions (0.001 M) to compensate for the negative charges of the phosphate backbone. Subsequently, the ionic concentration was brought to 0.1 M by the addition of KCl (mimicking physiological ionic strength)^30^. A cubic box was chosen with a total size of 109 Å to avoid RNA self-interactions in each direction for the simulations.

#### 2.3.2 Simulation Procedure

Initial structures were energy minimized using the steepest descent algorithm with a convergence threshold of 1000.0 kJ/mol/nm. Equilibration was then performed in the NVT ensemble and was carried out for 125 ps, followed by the NPT ensemble. Production simulation was carried out for 1 microsecond for each independent replicate. Hydrogen bond constraints were enforced throughout using the LINCS algorithm^31^. The md integrator^32^ was used for both equilibration and production phases. The system was maintained at a pressure of 1.0 bar during NPT using the C-rescale barostat^33^ with isotropic coupling, a relaxation time of 5.0 ps, and a compressibility of 4.5e-5, standard for water in GROMACS^16^. Temperature was controlled at 300 K using the v-rescale thermostat with separate coupling groups for solute and solvent, each with a coupling time constant of 1.0 ps. Electrostatics were handled using the Particle Mesh Ewald (PME) method, using standard Fourier spacing of 0.12nm. A Lennard-Jones cutoff of 0.9 nm was used for Van der Waals interactions, combined with dispersion corrections to energy and pressure to account for long-range interactions. The 0.9 nm cutoff was chosen to strike a balance between computational efficiency and the accuracy of simulating van der Waals interactions in a system, as has been done in other simulation studies of RNA systems^34^. Velocities were randomly assigned at the start of the NVT equilibration from a Maxwell-Boltzmann distribution at 300 K using a randomly generated seed. Five independent replicates were simulated for all systems, differing in their initial randomly distributed velocities.

### 2.4 Analysis Details

#### 2.4.1 Standard MD Analyses

We computed the radius of gyration, the atom-wise root-mean square deviation (RMSD), and the residue-wise root-mean-square fluctuations. We calculated the radius of gyration, a standard measure of molecular shape, using the GROMACS command *gmx gyrate*. We calculated the RMSD of the RNA with the GROMACS command *gmx rms* using the first frame of each simulation as the reference structure. We calculated the per-residue RMSF using the GROMACS command *gmx rmsf* to calculate the flexibility and mobility of individual residues.

#### 2.4.2 Center of Mass and sulfur-oxygen distance analysis

To assess the tightness of binding between ligands and targets, we calculated the center of mass (COM) distance from each ligand to the COM of the RNA binding site residues using the GROMACS command *gmx distance*. The thirteen binding site residues of the target were defined to be A6, U7, C8, A87, U88, G89 (members of the P1 loop), G11 (member of the J1/2 region of the internal L1 loop), and A45, A46, C47, U57, G58, C59 (members of the P3 loop), based on previous experimental studies^10^. We calculated the distance of the sulfur atom in SAM/SAH to the carbonyl oxygens of U7 and U88 using the ‘calc_bonds’ function from the MDAnalysis Python package^35,36,^ which calculates distances between any sets of atom pairs.

#### 2.4.3 Hydrogen bond analysis

Hydrogen bond analysis was carried out using the GROMACS command *gmx hbond*. We considered hydrogen bonds to be formed between a potential donor atom and a potential acceptor atom if the distance was less than or equal to the standard cutoff of 0.35 nm and the hydrogen-donor-acceptor angle was less than or equal to the standard cutoff of 30 degrees. We also characterized hydrogen bond occupancy of the *i*th bond *Oi*, according to the number of frames for which a particular h-bond was observed out of the total number of frames,

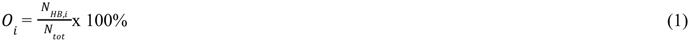

where *N_HB_*_,*i*_ is the number of frames in which bond is observed, and *N_tot_* is the total number of frames in the trajectory.

#### 2.4.4 pi-pi stacking analysis

To quantify π-π stacking interactions between the small molecule ligands containing aromatic rings (SAM and SAH) and RNA nucleotides throughout the molecular dynamics simulations, we adapted and modified the open-source Armadillo^37^ Python package built on the MDAnalysis package^35,36^. This code performs simple geometric calculations to identify T-stacked and parallel interactions between aromatic rings, and we expanded it to assess interactions over all frames in a trajectory. MD simulation data were loaded from TPR topology and XTC trajectory files with water molecules stripped out, with analysis restricted to the atoms corresponding only to the RNA-ligand complex. Atoms forming aromatic rings were identified separately for the ligand (SAM and SAH residues) and thirteen RNA-binding site residues using residue-specific atom name selections. For each frame of the trajectory, intermolecular π-π stacking interactions were identified between ligand and RNA rings using geometric criteria: inter-ring distances between 2.0-5.0 Å and angular classifications based on the angle (α) between ring normal vectors. Interactions were categorized as parallel-displaced (α < 45°) or T-shaped (α > 80°) configurations.

#### 2.4.5 Construction of dynamical correlation network

To better assess the effects of the binding of the ligand to the overall riboswitch complex, we perform a dynamic correlation network analysis using the *dynetan* Python package^38^. In brief, all RNA residues and the ligand are mapped to nodes in a network whose edges are weighted by the correlations between the atoms assigned to each node over the course of the simulation. Correlation in the dynetan package is calculated with a generalized correlation coefficient that forms an approximation to the mutual information between two nodes and provides a measure of whether the two are moving together. Analyzing the resulting network provides information about more global motions of the complex and permits us to better assess the effect of the ligand and its important contacts in context.

For the network analysis, following the standard practice laid down by the authors of the *dynetan* package, we represent each RNA residue as two nodes, one representing the phosphate backbone and one representing the sugar ring(s). We represent the SAM and SAH ligands as three nodes apiece, representing respectively the adenosine ring, the methionine tail, and the central ribose ring and sulfur atom, as these are approximately our three regions of interest. We divide JS4 into five nodes to keep the nodes roughly the same size as in the other ligands and residues (about 6-11 atoms per node). All nodes are represented by a central atom: for RNA residues, we employ the phosphorus atom and the joining nitrogen, as described in the *dynetan* tutorial; for the SAM and SAH ligands, the nodes are centered at a similar nitrogen to adenine, at one of two central carbons in the tail, and at the sulfur atom which we expect to be of importance to ligand binding. For JS4, we determine node centers based on spatial spread and identifying those atoms that are highlighted as participating in particular hydrogen-bonding interactions (**section 3.3**). (See **Fig. S1** for an illustration of atom-to-node mappings for the RNA residues and the ligands.) To construct dynamic correlation networks between the nodes, we employ five windows corresponding to the five independent replicates and use a cutoff distance of 4.5 angstroms and a contact persistence cutoff of 75%. In other words, nodes are only considered to be in contact when they are less than 4.5 angstroms apart for more than 75% of the trajectory frames. Although the resultant network can depend quite strongly on these parameter choices, because we are performing a comparative calculation between different systems, we expect this to be sufficient to assess system trends. In addition, we do not calculate contacts between nodes in the same residues or consecutive residues. Cartesian calculations are performed using all 10,0001 frames in each of the five independent replicates.

#### 2.4.6 Network analysis on dynamical correlation networks

Once networks have been constructed representing the dynamical correlations of each window, we perform a set of standard network analyses using a combination of the *dynetan* package and the *networkx* Python package^39^, which provides direct access to network calculations.

#### Community detection

To understand which parts of the protein move together, we perform a community analysis with the Dynetan package, which employs a heuristic algorithm^40^ instantiated in NetworkX (as community.best_partition) to optimize community modularity according to node assignment to communities. Modularity is essentially a measure of the “structure” of a network. High modularity implies that there are strong (highly weighted) edges between nodes within communities, while edges between nodes that are in different communities are weak. It is defined as,

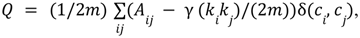

where *m* is the sum of all edge weight, *A* is the adjacency matrix of the graph, *ki* is the weighted degree of node *i*, γ is a resolution parameter, and δ(*c_i_*, *c_j_*) is the Kroneker delta function, which is 1 if *i* and *j* are in the same community, and zero otherwise. For each replicate, we calculate the best partition of the associated network, and of these five partitions, we identify the one with the highest modularity as the reference community. Then the communities associated with the different replicates are assigned based on their overlap with the reference community. We do not consider any communities with fewer than 10 nodes in the reference.

#### Eigenvector Centrality

To identify which parts of the ligand-riboswitch complex exert the greatest influence over its motions, we compute the eigenvector centrality of each node in the network for each network associated with a different system and replicate using the NetworkX function ‘eigenvector_centrality’; then we report the average eigenvector centrality per node for each system and its standard deviation. Eigenvector centrality for a node *i* is the *i*th entry in the right eigenvector of the adjacency matrix of the graph corresponding to the largest eigenvalue. This is the proportion of dynamical “information” that will be retained on the node after a long time; it can also be thought of as a generalization of degree centrality, from computing the importance of a node based on the connections of its nearest neighbors to computing the importance of the node based on the connections of the neighbors of the neighbors in the limit of infinite neighbors^41^.

#### Edge Betweenness

To identify those pairs of nodes through which the greatest amount of information flows, we further compute the edge betweenness centrality of the edges in each network associated with a different system and replicate using the built-in Dynetan package computation of edge betweenness, which is a wrapper for the NetworkX function ‘edge_betweenness_centrality’; then we report the average edge betweenness centrality per node for each system and its standard deviation. Edge betweenness centrality is a different way to assess the flow of information in a network: it is computed as the sum of all-pairs shortest paths that include the edge in question^42^,

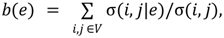

where *e* is the given edge, *V* is the set of nodes (vertices) in the graph, and σ(*i*, *j*) denotes the shortest path between nodes *i* and *j*. This functions as a very different metric for assessing importance, as it demonstrates the edges that, if cut, would force the most rerouting in the network, showing which edges are important to maintain the dynamical motions of the riboswitch complex under each set of conditions.

## 3. RESULTS

### 3.1 Virtual screening identifies a potential binder of the SAM-I riboswitch

We performed molecular docking to produce initial conformations for MD simulation and to investigate the type of binders it could produce from a repurposed list of general RNA binders. Once identified, our study sought to enhance the understanding of how the top-ranked ligand (JS4), along with SAM and SAH (chemical structures shown in **Fig. S2** along with atom naming schema), might bind and interact with the SAM-I riboswitch RNA target, providing insights into its likely efficacy and selectivity as therapeutic agents as well as design constraints for future research.

We re-docked the co-crystallized SAM ligand into the SAM-I riboswitch target’s binding site to validate the accuracy of the FITTED molecular docking software for our application and to produce an initial docked pose for later simulation. We did five runs using the SAM-I riboswitch RNA target and its co-crystallized ligand, SAM. We calculated the RMSD of the final five poses of the ligand from its pose in the crystal structure. Among the five runs, the most favourable **(see Fig. 2a)** possessed an RMSD of 1.22 Å (see **Table S1** in the Supplementary information for a full list of predicted binding energies). Indeed, all poses possessed RMSDs of ≤ 2.0 Å to the experimental pose, which is considered good, especially for RNA-ligand interactions, where flexibility and conformational changes can impact docking precision^43,44^. The RMSD value produced in our study is indicative of a trustworthy level of agreement between the docked pose and the experimentally crystallized conformation. The average predicted binding energy was -103+/-4 kCal/mol, suggesting irreversible binding. We next docked the negative control (SAH) into the target, predicting an average binding energy of 95 +/- 9 kCal/mol, suggesting potentially reversible binding, and producing a similar initial pose to that of SAM (Fig. **2b**).

**Figure 2:**
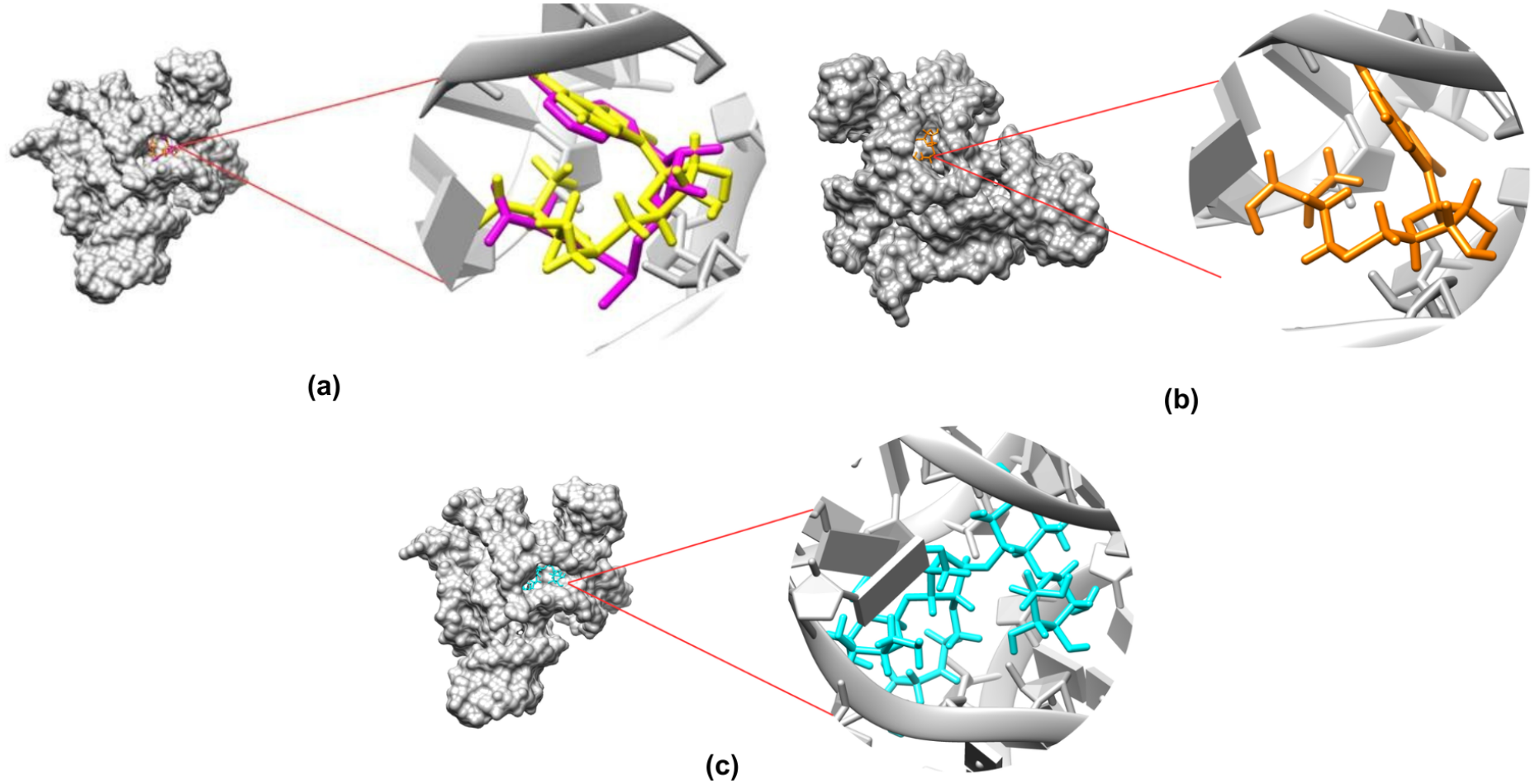
Docking poses for initial simulation. (a) The best SAM pose (yellow) from self-docking with an RMSD of 1.22 Å from the experimental SAM pose (magenta), (b) most favourable docked SAH pose (orange) and (c) most favorable docked JS4 pose (cyan). Docking performed with FITTED molecular docking software. Images rendered using UCSF Chimera^45^.

Finally, we docked our curated 150 small molecules to our RNA target. We identified JS4 as the top binder (**see Fig. 2c**) to our RNA target with an average predicted binding energy of -140 +/- 10 Kcal/mol (see **Table S3** in the Supplementary information for a full list of predicted binding energies). The lower binding energy predicted for JS4 than for SAH or SAM suggests the potential of irreversible binding; however the number should be interpreted with caution, as docking softwares are still less accurate for RNA than for protein binding. This underscores the need for a more comprehensive analysis to understand potential ligand binding.

### 3.2 SAM and JS4 are more stably bound than SAH

From calculations of the radius of gyration and the RMSD, we ensured that all complexes were equilibrated and reasonably stable over the course of the simulations (see Section S2). Next, to assess the strength and stability of ligand binding, we calculated the center of mass (COM) distance between the ligand and the binding site of SAM on the SAM-I riboswitch (**Fig. 3**). (See also **Figs S2**-**3** for individual replicate distances and further representative snapshots.) The native ligand remains tightly and stably bound throughout all simulations, maintaining a COM distance of about 0.1 nm regardless of system initialization (**Fig. 3a**, yellow and pink traces). For the SAH complex, we observe a gradual increase in COM distance from ∼0.2 nm to ∼0.6-0.7 nm, indicative of partial dissociation in several of the replicate simulations. While the COM of the larger JS4 sits further from the center of the SAM binding site (at ∼0.6 nm), the trace does not change across the course of the simulation, indicating also a differing but still stable binding mode. Thus, SAM and JS4 do demonstrate irreversible binding in simulation, while SAH is less stable, which bears out the results of docking at a qualitative level.

**Figure 3:**
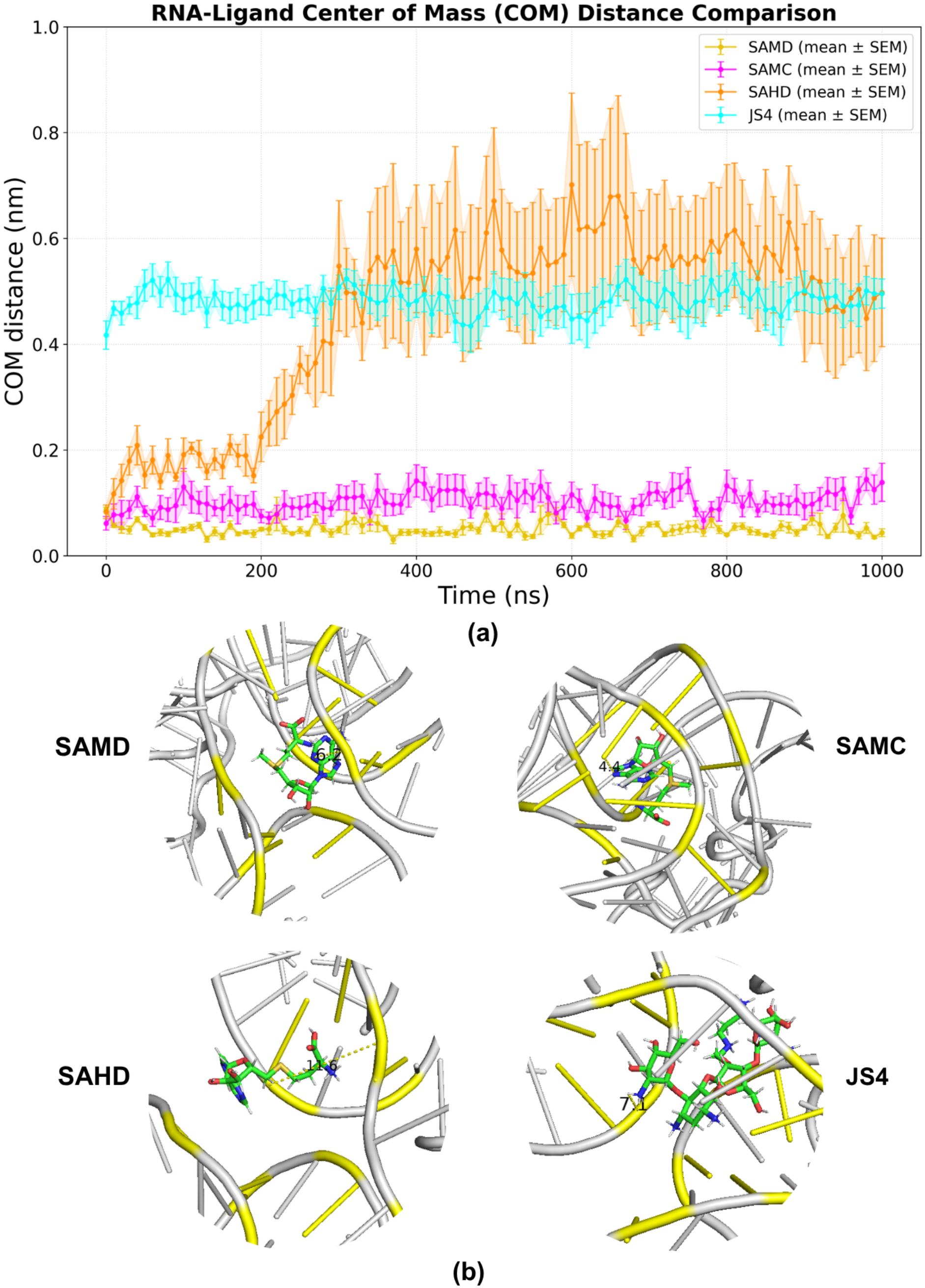
COM distance between ligand and SAM binding site vs time for (a) SAM ligand docked (yellow) with RNA, SAM ligand crystallized (magenta) with RNA, SAH ligand docked (orange) with RNA, JS4 ligand docked (cyan) with RNA. Error bars represent the standard error taken across five independent replicates. (b) Illustrative snapshots of MD simulations at 500 ns. Snapshots rendered with PyMOL^11^.

### 3.3 Analysis of hydrogen bonds formed between riboswitch and ligands suggests that JS4 interacts more strongly than SAH with similar residues, but less strongly than SAM

To assess the molecular-level interactions controlling RNA-ligand behaviour, we calculate the number of hydrogen bonds between RNA and the ligand (see **Fig. 4**), as well as the occupancies of individual hydrogen bonds with binding site residues as a measure of their persistence over time (see **Fig. 5**, error in **Fig S5**). JS4, SAMD and SAMC maintain a higher number of hydrogen bonds on average in all five replicates (cf **Table S6** in the SI) than SAH. The RNA-SAH complex shows the lowest number, dropping from ∼6-7 to ∼2-4 hydrogen bonds over the course of the simulation timescale, concomitant with the looser binding mode also reflected with the increasing COM distance.

**Figure 4:**
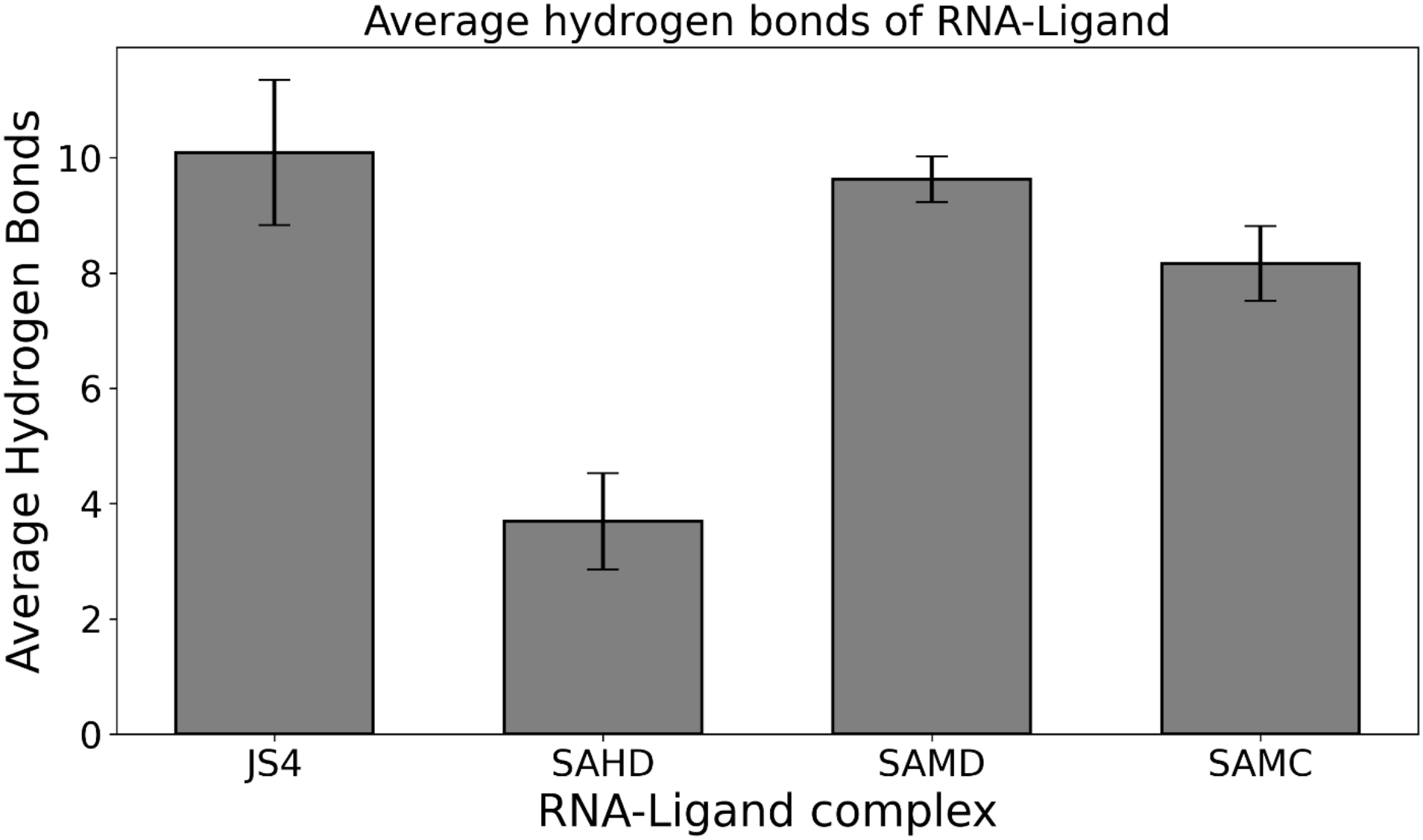
Average number of hydrogen bonds in four RNA-ligand complexes

**Figure 5:**
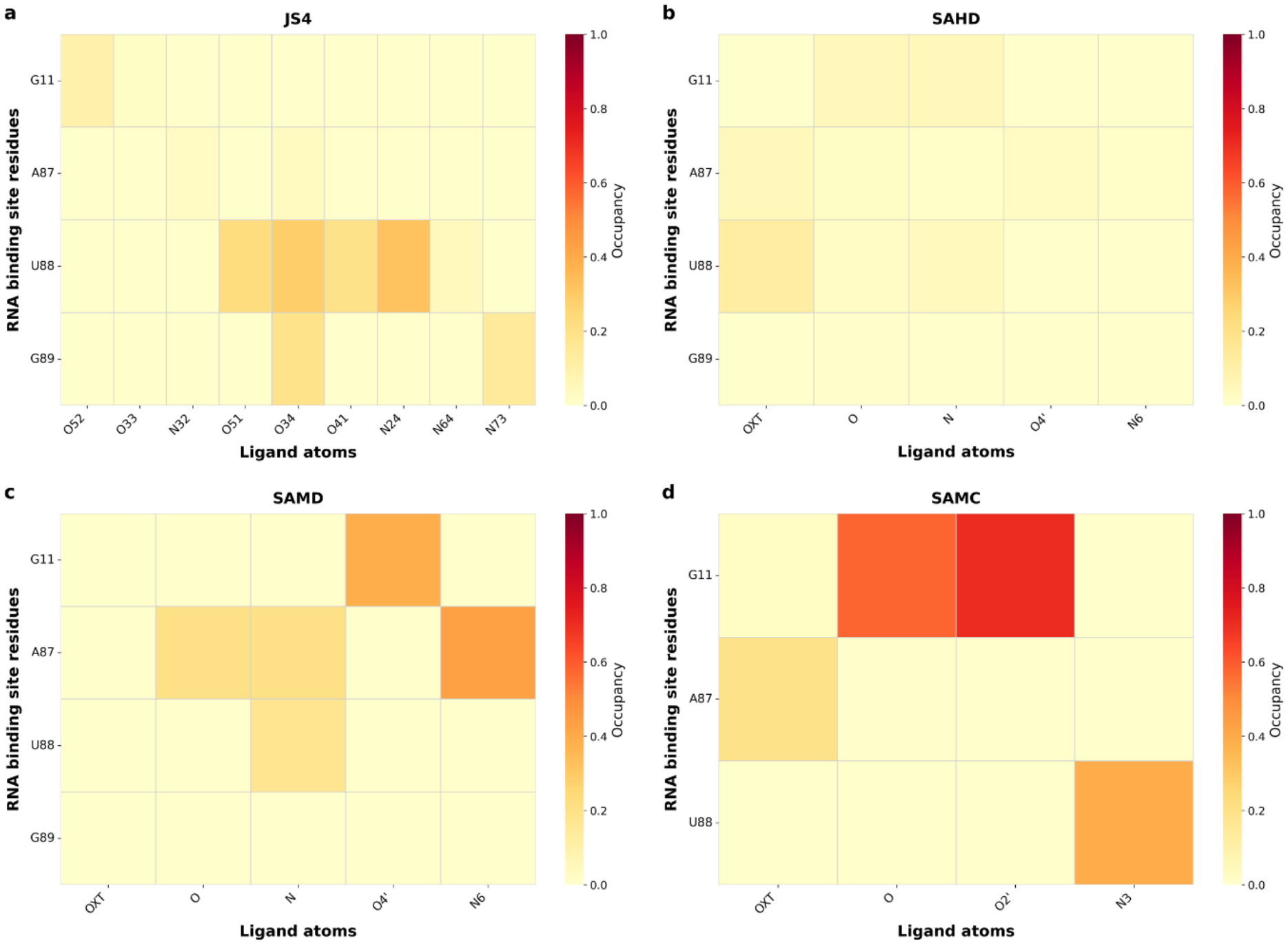
Persistence of hydrogen bonds formed between ligand atoms and RNA binding site residues (a) RNA-JS4 complex (docked) (b) RNA-SAH complex (docked) (c) RNA-SAM complex (docked) (d) RNA-SAM complex (co-crystallized).

Of the thirteen RNA residues defined as part of the binding site, four are responsible for all hydrogen bonding interactions with the ligands in our simulations: residues G11, A87, U88, and G89 (cf. **Fig. 1c**). SAM (docked) shows moderately persistent interactions with G11 at occupancy ∼0.4-0.5 where the adenosyl sugar ring forms a contact with G11 residue at the O4’ ribose ligand atom (**Fig. 5c, Fig. S2** showing ligand atom labeling). It also demonstrates moderately persistent interactions of the adenine base with residue A87 (atom N6), and weaker interactions of the methionine tail with the same residue (atoms O, N, persistence ∼0.2). Similarly, the methionine tail interacts weakly with U88 (atom N, persistence ∼0.2). SAM (crystallized) possesses similar interactions with the same three residues (**Fig. 5d)**; however, its h-bonding with G11 and U88 is more persistent than in the SAMD system, and the interactions (with >0.8 persistence) occur with the O2’ atom of the adenosyl ring, and with the O atom of the folded-over methionine tail (∼0.6 persistence). U88 bonds with the adenine base (N3 ligand atom, ∼0.4-0.6 occupancy). It has fewer and more transient interactions with residue A87, mediated solely by the methionine tail. For the SAH system, (**Fig. 5b**), interactions are substantially more transient, but primarily with the same three residues. The homocysteine tail (at atoms O and N) forms transient interactions (forming a hydrogen bond under ∼10% of the time) and slightly more persistent interactions with U88 (atom OXT, occupancy around ∼0.2 and atom N, <0.1). We observe very transient interactions between SAH and A87. Similar to SAH, JS4 has more substantial interactions with U88, though its interactions are more persistent than those of SAH (multiple atoms forming h-bonds 40-50% of the time), as well as transient interactions with residues G11 and G89 (∼0.2 occupancy or a little higher for 2 atoms) and very transient interactions with A87 (**Fig. 5a)**.

### 3.4 Simulations highlight pi-pi stacking interactions with riboswitch residue C47

To more fully characterize the non-covalent molecular-level interactions between the SAM-I riboswitch and SAM and SAH, we investigate the contribution of π-stacking to the stability of ligand binding, since it is a major contributor to the stability of RNA-ligand complexes^46^. In our simulations, the great preponderance of geometrically-defined pi-pi stacking interactions occurs between the ligand and residue C47 for all ligand systems containing aromatic rings (**Table S7, Fig. 6**), with only transient or replica-specific interactions formed with other residues. In the SAMC system, we observe that the ligand and residue C47 maintain a geometry concordant with parallel pi-pi stacking at an occupancy of 99 ± 0.5%. In the SAMD system, we observe a similar parallel-stacking geometry but at only 46 ± 1% occupancy, likely due to the greater flexibility of the ligand also illustrated by its more promiscuous h-bonding behavior. On the contrary, SAH exhibits parallel pi-pi stacking geometry with C47 only transiently (10 ± 4%). Overall, this suggests the importance of pi-pi stacking with residue C47 to help maintain tight binding.

**Figure 6:**
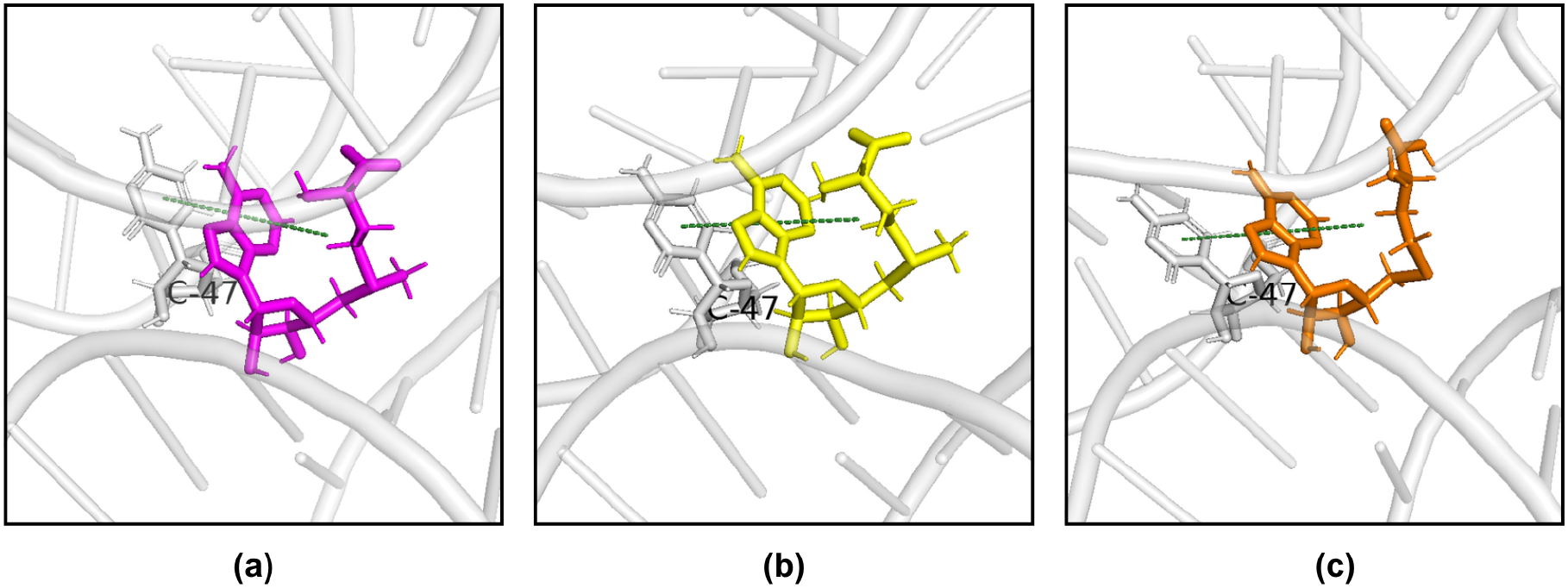
Molecular dynamics simulation snapshots showing π-stacking interactions between the aromatic adenosine ring systems of three ligands and the cytosine 47 (C47) base of the RNA binding site.The gray cartoon represents the RNA backbone and surrounding residues; the colored stick representations show the ligand-binding pose. Green dashed lines indicate π-π stacking interactions between the aromatic ring systems. (a) π-stacking in SAMC (magenta) with C47 residue of RNA, (b) π-stacking in SAMD (yellow) with C47 residue of RNA, (c) π-stacking in SAHD (orange) with C47 residue of RNA. Snapshots rendered with PyMOL^11^.

### 3.5 Residues U7/U88 of the SAM-I riboswitch remain in dynamic proximity to sulfonium ion of SAM

To complement previous experimental observations which reported the importance of binding between the sulfonium ion of SAM and residues U7/U88 in the riboswitch, we calculated the average distance from the sulfonium ion in SAM and the sulfur atom in SAH to the carbonyl O2 oxygens of U7/U88 in our simulations (**Fig. 7**). In the SAMD complex **(Fig. 7a)**, we observed a largely unchanging distance between the sulfonium and both oxygens over time, with the sulfonium remaining at an average of *r _U7_* ∼3. 8 angstroms from the U7 oxygen and an average of *r_U88_* ∼5 angstroms from the U88 oxygen. Assuming an electrostatic interaction between these atoms with similar charges on both oxygens, the interaction strength scales inversely with the square of the distance. This yields a ratio of (*r_U_*_88_/*r _U_*_7_)^2^ ≈ 1. 7, indicating ∼1.7 times stronger interaction with U7 than with U88, while both interactions remain steady and persistent. In the SAMC complex (**Fig. 7b**), we note a similar interaction pattern, though the distances to both U7 and U88 are slightly greater, with average distances of 4.73 ± 1.05 Å between the sulfonium and U7 and 5.72 ± 0.89 Å between the sulfonium and U88. This yields a ratio of (*r_U_*_88_/*r U*7)^2^ ≈ 1. 5, again favouring U7. Conversely, in SAH, (**Fig. 7c**) we observe generally greater distances, and there is more of a spread between different replicates, with replicate 3 maintaining a distance of about 4-6 Angstroms with both oxygens, while replicates 1 and 4 remain about 6 Angstroms away from U88, but drift to about 7-8 Angstroms away from U7, and replicates 2 and 5 fluctuating to distances of 10-15 Angstroms from U88 and U7. Just from distance considerations, this would lead to an average interaction strength drop of at least fifty percent, but, of course, there is a substantial drop as well in the (partial) charge on the sulfur. Overall, similar to experiment, we observe persistent interactions between the sulfonium and both U7 and U88, while there are far less persistent interactions between those residues and the neutral sulfur of SAH. This does indicate the role of the charge-charge interactions with U7 and U88 in stabilizing the ligand binding, supported by h-bonding and pi-pi stacking with other residues.

**Figure 7:**
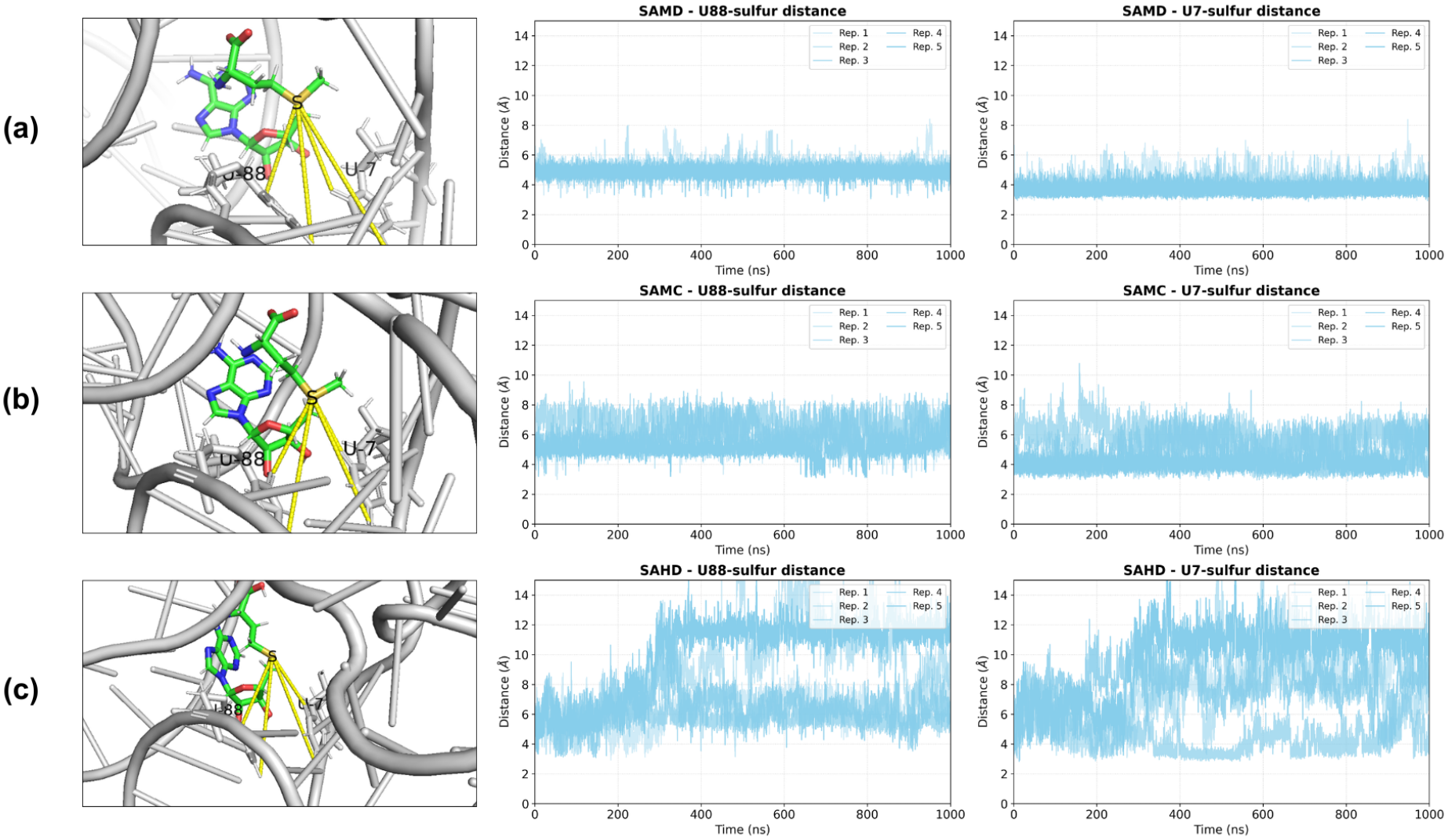
Distances between sulfur (of SAM and SAH) and U7/U88 (O2) (of RNA) along with MD snapshots distance of sulfur and U88/U7 RNA residues shown with yellow line. (a) SAMD-RNA complex, (b) SAMC-RNA complex, (c) SAHD-RNA complex. Snapshot rendered with PyMOL^11^.

### 3.6 SAM binding decreases flexibility in the P1 and P3 loop regions

To assess our hypothesis that SAM binding alters flexibility in the binding loops, we calculated the root-mean-square fluctuations of the RNA and how it varied across relevant regions during the course of the simulations (**Fig. 8**): the binding site, (panel A), the P1 loop (panel B), the P3 loop (panel C), and all other residues (Panel D). (In **Fig. S6**, we also present the residue-wise RMSF.)

**Figure 8:**
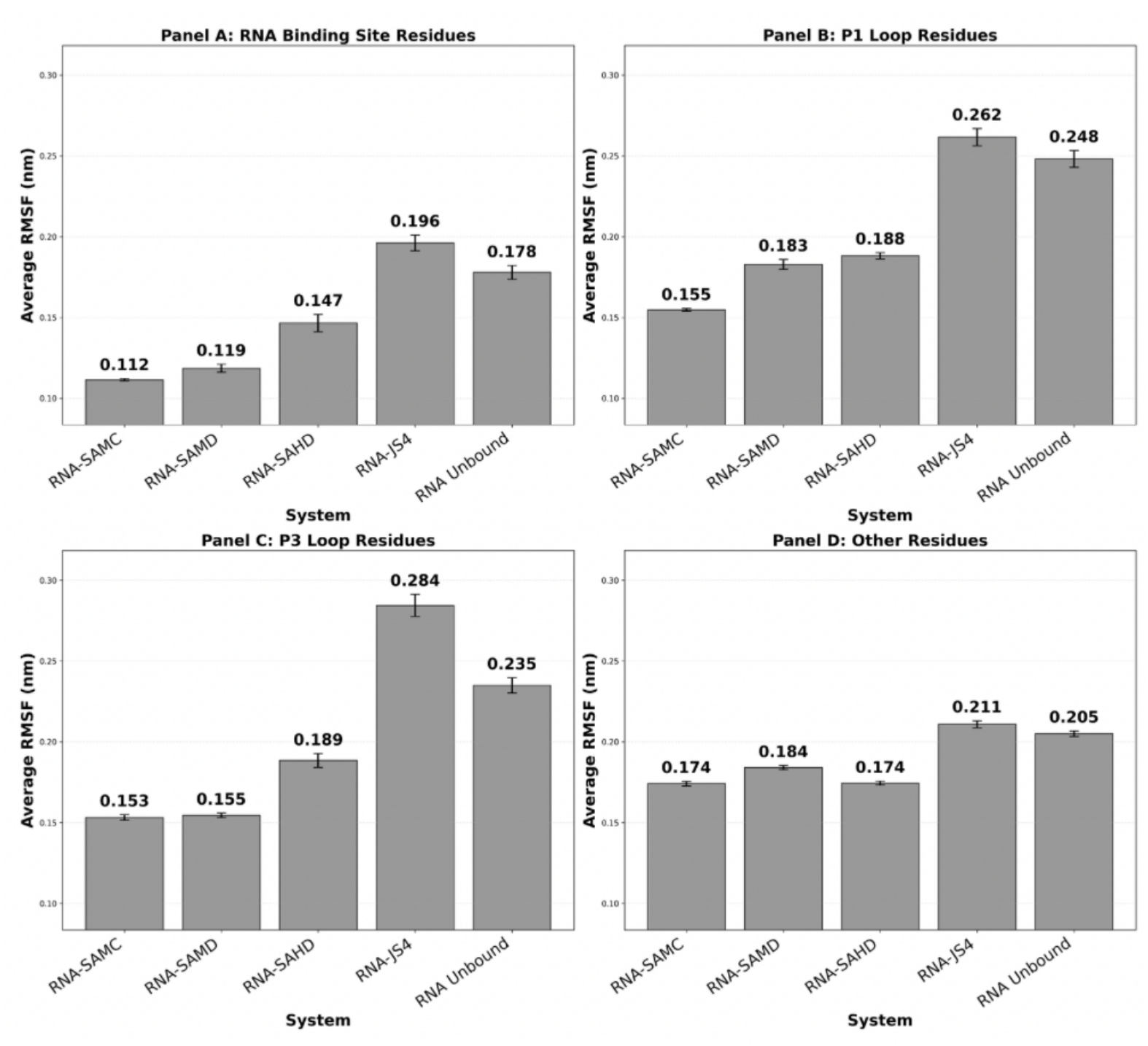
Bar plots presenting average RMSF values across distinct RNA regions: Panel A, binding site residues; Panel B, P1 loop residues; Panel C, P3 loop residues; and Panel D, all other residues.

In terms of the binding site residues, SAMC demonstrates the lowest flexibility (average 0.112 nm), with SAMD a close second (average 0.119 nm). To our surprise, the SAH system was next (average 0.147 nm), with the unbound system (average 0.178 nm) and the JS4 system (average 0.196 nm) displaying higher flexibility in the binding site **(Fig. 8a)**. In other words, all ligand-bound systems except JS4 demonstrate reduced flexibility at the binding site compared to unbound RNA, with the SAM ligand providing the most stabilizing effect.

The P1 loop region exhibits higher flexibility across all systems compared to the binding site **(Fig. 8b)**. Notably, RNA-JS4 displays particularly elevated flexibility (average 0.262 nm), the highest among all ligand-bound states, and higher even than the unbound state (average 0.248 nm), suggesting that JS4 binding may permit or even induce significant P1 loop conformational changes compared to the native bound structure. In addition, we note that the SAMC residues show the lowest flexibility in the P1 loop residues (average 0.155 nm) compared to SAMD or SAH (average 0.183 nm and 0.188 nm, respectively).

The P3 loop residues **(Fig. 8c)** display striking ligand-dependent flexibility patterns, with SAM-bound systems maintaining the lowest flexibility (SAMC: 0.153 nm, SAMD: 0.155 nm), intermediate flexibility in the SAH system (0.189 nm), elevated flexibility in unbound RNA (0.235 nm), and markedly elevated flexibility in JS4-bound RNA (0.284 nm)- the highest among all systems. This indicates that despite JS4’s persistent interactions with residues near the active site, it fails to stabilize the P3 loop.

In residues outside the binding site, P1 loop, and P3 loop **(Fig. 8d)**, we observe intermediate and relatively uniform flexibility across all systems. RNA-SAMD (average 0.184 nm), RNA-SAMC (average 0.174 nm), and RNA-SAHD (average 0.174 nm) all display similar flexibility. RNA-JS4 (average 0.211 nm) and RNA-Unbound (average 0.205 nm) show moderately higher flexibility, with JS4 again demonstrating the highest RMSF. The comparatively small differences between ligand-bound and unbound complexes in these uninvolved regions indicate that ligand binding primarily affects local structure at the binding site, P1 loop, and P3 loop rather than inducing significant global conformational changes throughout the RNA structure.

Considering the average RMSF of “other residues” as setting a standard level of fluctuations for the system, we may also compare the fluctuations of the binding site and P1 loop with respect to this reference. For the binding site, SAM displays the greatest comparative suppression, with ∼65% as much flexibility in the binding site as in other residues for both crystallized and docked complexes; SAH and unbound are commensurate at 84% and 86%, respectively, and JS4 displays little suppression in flexibility at the binding site (93% flexibility compared to the uninvolved sites). For the P3 loop, SAM shows moderate suppression (∼84-87% of other residues), while SAH shows a small enhancement (∼109%) and JS4 shows an appreciable one (∼122%), indicating heightened flexibility greater even than the unbound state (which only has an enhancement of ∼114%). The pattern for the P1 loop differs slightly, with SAMD fluctuating about as much as the other sites on average (99% RMSF in the P1 loop residues as compared to the other residues), and SAMC showing a small amount of suppression in P1 loop motion (89% RMSF in the P1 loop residues as compared to the other residues). SAH displays slightly heightened flexibility (108%), and JS4 and unbound display moderately heightened flexibility (124% and 121%) in the P1 loop compared to the other states.

Overall, our simulations show binding site residues which tend to display lower flexibility than the other residues, though to a negligible extent when bound to JS4. SAM suppresses fluctuations of the P1 and P3 loops in simulation, while SAH tends to maintain them and JS4 and the unbound state are associated with enhanced flexibility in the region.

### 3.7 Network analysis suggests that the SAM ligand modulates coordinated motion of the P1 and P3 loops

In this section, we delve more deeply into the predicted effects of the ligand on global fluctuations through the calculation of dynamic correlation networks and an accompanying network-based analysis. Such analyses were originally developed to better describe allosteric motions in proteins, but the same theory holds for RNA. Indeed, because RNA tends to be more flexible than proteins, we propose that this form of analysis is even more valuable for RNA simulations which may be less likely to show all the large-scale motions of which they are capable during accessible simulation timescales.

In **Fig. 9**, we show how the nodes of the network divide into communities, neglecting those with under ten nodes. In this context, a community is essentially a portion of not-necessarily-proximal parts of the complex that move together and do not have strong correlated motions with parts of the complex belonging to other communities. We note that there are differences in the number of nodes participating in large (>10 nodes) communities depending on the system. Both SAM-bound systems have larger communities formed by greater numbers of participating nodes than the others (112 nodes participating in large communities/187 nodes in the network = 60% for SAMC and 92/181=51% SAMD) compared to ∼40% for the other three (72/183 for SAH, 74/185 for JS4, 75/182 for the unbound state). Thus, there is evidence of more global communication in the simulations containing the native ligand, even though the total number of large communities is similar in all cases (8-9). Concomitantly, the SAH ligand nodes do not participate in any large communities, and only one out of five of the JS4 ligand nodes does, compared to all three SAM ligand nodes. In addition, the native ligand systems maintain a larger and more consistent community containing two of the ligand nodes (those associated with the two rings, including the sulfur atom) and many of the binding site residues from both the P1 and P3 loops (assigned ID 1.0 in **Fig. 9a** and **9b**, splitting into two subcommunities with IDs 1.0 and 1.1 in replicates 3 and 5 of **9b**). Specifically, consider the community in which the binding site residue U88 takes part, which we shall call C-U88. In the SAMC system, C-U88 contains the binding site residues A6, U7, A87, U88 (by definition), as well as G89 of the P1 loop, C47, U57, and G58 and usually A46 of the P3 loop. It also contains the non-binding site residues C48 and G56 of the P3 loop most of the time. Although there is slightly more replica-specific variation in SAMD, the same community is generally observed, albeit sometimes also containing the binding site residue C8 of the P1 loop and the non binding site residue A10 of the J12 loop. The primary difference in SAH, JS4, and the UB system is the loss of the P3 loop residues from this community, including C47, and, primarily in the UB system, the addition of most of the rest of the P1 loop residues.

**Fig. 9.**
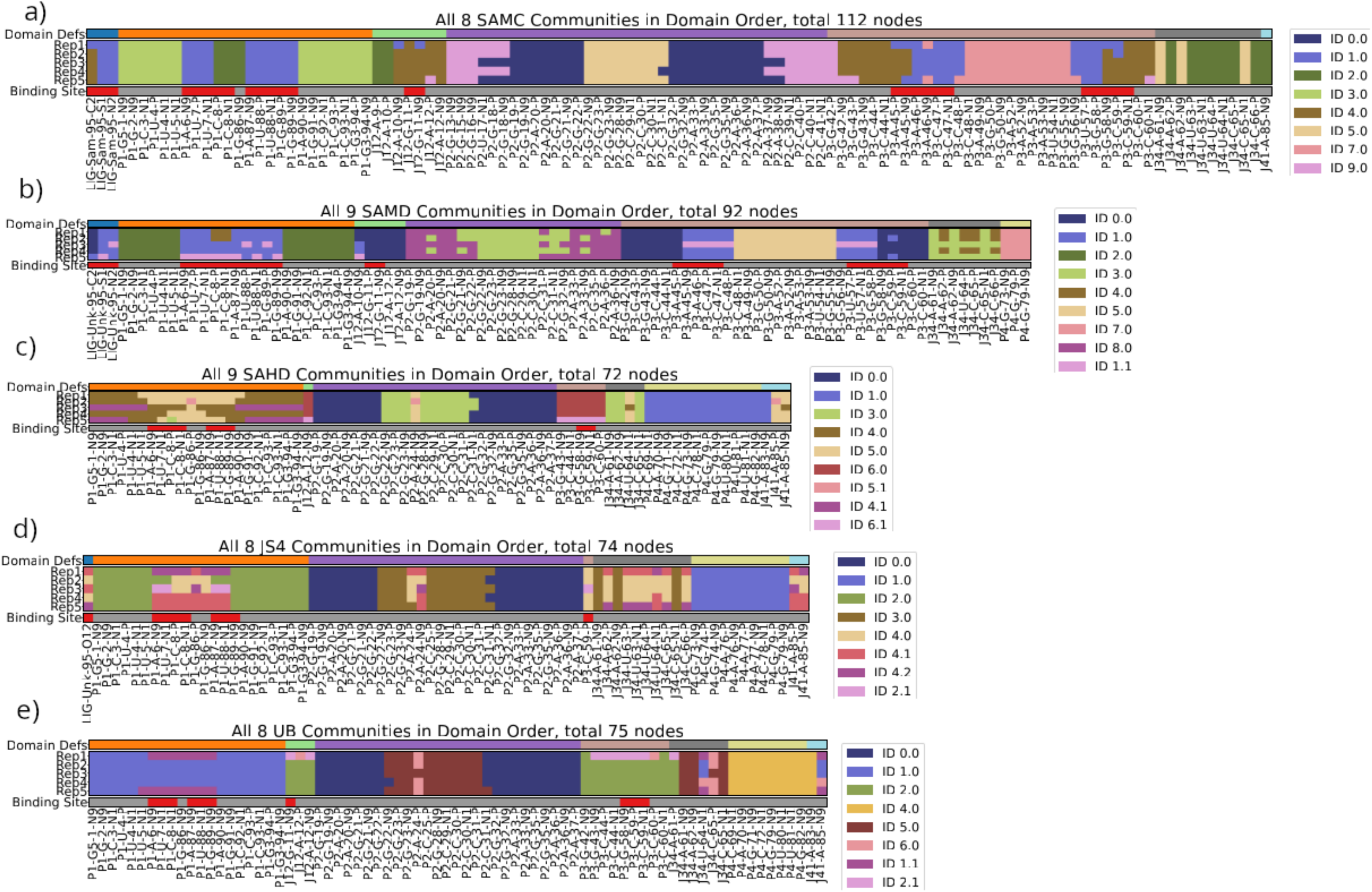
Major communities in dynamical correlation networks of riboswitch-ligand complexes for a) SAMC system, b) SAMD system, c) SAHD system, d) JS4 system, e) unbound system. The horizontal axis lists all nodes that participate in communities with greater than 10 members. Each node is colored by (top strip) the domain to which it is assigned, (bottom strip) whether it is part of the experimentally-determined binding site (including the ligand), and (middle five) the community to which it is assigned in each of the replicates. Color maps differ between systems for the community labeling. Color map for the top strip is as follows: blue—ligand, orange—P1 domain; light green—J12 region of the L1 internal loop; purple—P2 domain; light brown—P3 domain; dark grey—J34 region of the L1 internal loop; yellow—P4 loop; light blue—J41 region of the L1 internal loop. On the bottom strip, the binding site is highlighted in red; all other residues are colored grey. Community IDs and colors are assigned arbitrarily; however decimals indicate subcommunities that are separate in one replicate but part of the same community in the reference replicate (i.e. if the nodes of community assigned ID 1.0 in the reference network belong primarily to two communities in a different replicate, those two communities will be assigned IDs 1.0 and 1.1 in that replicate.)

Although the SAM tail node participates in C-U88 in replica 1 of the SAMC system and replica 5 of the SAMD system, it primarily belongs rather to the same community as residue G11 of the binding site/J12 loop (assigned ID 4.0 in SAMC, 0.0 in SAMD). The community containing residue G11 of the J12 loop (referred to as C-G11) appears similar across SAMC, SAMD, and UB systems. Specifically, in addition to G11, in SAMC and SAMD, C-G11 primarily contains binding site residues A45, G58, and C59 of the P3 loop, and non-binding site residues A10 and A12 of the J12 loop, G42, G43, C44, and C60 of the P3 loop, and A61 of the J34 loop. In UB the story is similar, except that binding site residue A45 and non binding site residue A10 are not part of the community.

In terms of ligand participation in large communities, the JS4 system does demonstrate a community (assigned ID 4.0 in **Fig. 9d**) containing a single ligand node and several of the binding site nodes, but only in replicate 2 is there participation of the ligand and binding site nodes from both the P1 and P3 loops in the same community. In all other replicates, these split into sub-communities in different patterns. Meanwhile, not only do the ligand nodes in the SAH system not participate in large communities, the binding site residues of P1 and P3 do not participate in shared communities, as is also the case for the unbound system. Additionally, we observe differential P3 loop involvement in large communities across different systems. The P3 loop forms ∼30% of the nodes participating in large communities for SAMC and SAMD (31/112, 31/92 nodes respectively; light brown in the top strips of **Fig. 9**) but only ∼10% for the unbound and SAH systems (9/75, 5/72 nodes respectively) and is almost entirely absent in the JS4 system (only 1 node of the 74 nodes associated with communities of >10 members is associated with the P3 loop—one of the nodes representing residue C59). We also observe differential P4 loop involvement in large communities (light yellow-green in the top strips of **Fig. 9**)—there is little-to-no involvement for native communities but fairly substantial involvement in the unbound, SAH, and JS4 systems, where it tends to form a community by itself (assigned ID 1.0 in **Fig. 9c,d**, and ID 4.0 in **Fig. 9e**).

In **Fig. 10** we present the average edge betweenness for high betweenness edges. (For clarity, in **Fig. S7** we show the average edge betweenness for all edges with non-zero betweenness, and in **Fig. S8** we qualitatively visualize the edge betweenness weight mapped onto the three-dimensional complex.) Edge betweenness is defined as the normalized number of shortest paths for which that edge is a component. In the context of a dynamical network analysis, high betweenness implies that there is a strong dynamical connection between the nodes in question and that they form a “bridge” carrying a large proportion of the dynamical information of the network. We might think of regions of high betweenness as regions through which much of the dynamical information “flows.” For both native ligand-bound complexes (**Fig. 10a,b**), we observe high-betweenness edges involving the sulfur atom of the SAM ligand (C8-Sulfur edge in both, G89-Sulfur edge in both, and U88-Sulfur edge in the SAMD system). Conversely, none of the high-betweenness edges of the SAHD system include the ligand, despite there being a higher number of edges with betweenness greater than 0.6 (see also **Fig. S8** where it is evident that the shortest paths of the network flow *around* the ligand). The highest betweenness edges occur within the P2 loop (orange bars), e.g. G22-C30. In the JS4 system, there are four high-betweenness edges involving the ligand, but they occur in different regions from those in the SAM systems: in the SAM systems, betweenness is concentrated in the region where the ligand contacts the P1 binding site and the P1 binding site encounters the J34 loop (high betweenness edge linking C8-U64), with some moderate betweenness edges in the P2 and P3 loop regions, as well as one (G28-C66) connecting the P2 loop with the J34 loop. Conversely, in the JS4 system, the highest betweenness is in the P2 loop (G22-C30 edge, similar to SAH) and the J34 loop (A62-C65). The high betweenness edges involving JS4 occur in different regions to those involving SAM: JS4-P3 (C59-JS4, C60-JS4) and JS4-J34 (involving U64 and C65-related nodes). (See also **Fig. S8** where the betweenness in the JS4 system tends to be a little higher, and less commonly reaches the bottom of the P3 loop). We further note that the unbound system tends to have lower overall betweenness, with almost all of its high betweenness edges being in the distal P2 loop (see also **Fig. S7**), suggesting less overall correlated motion for this system than the others. The P4 loop remains uninvolved in transfer of dynamical information for all simulated systems.

**Fig 10.**
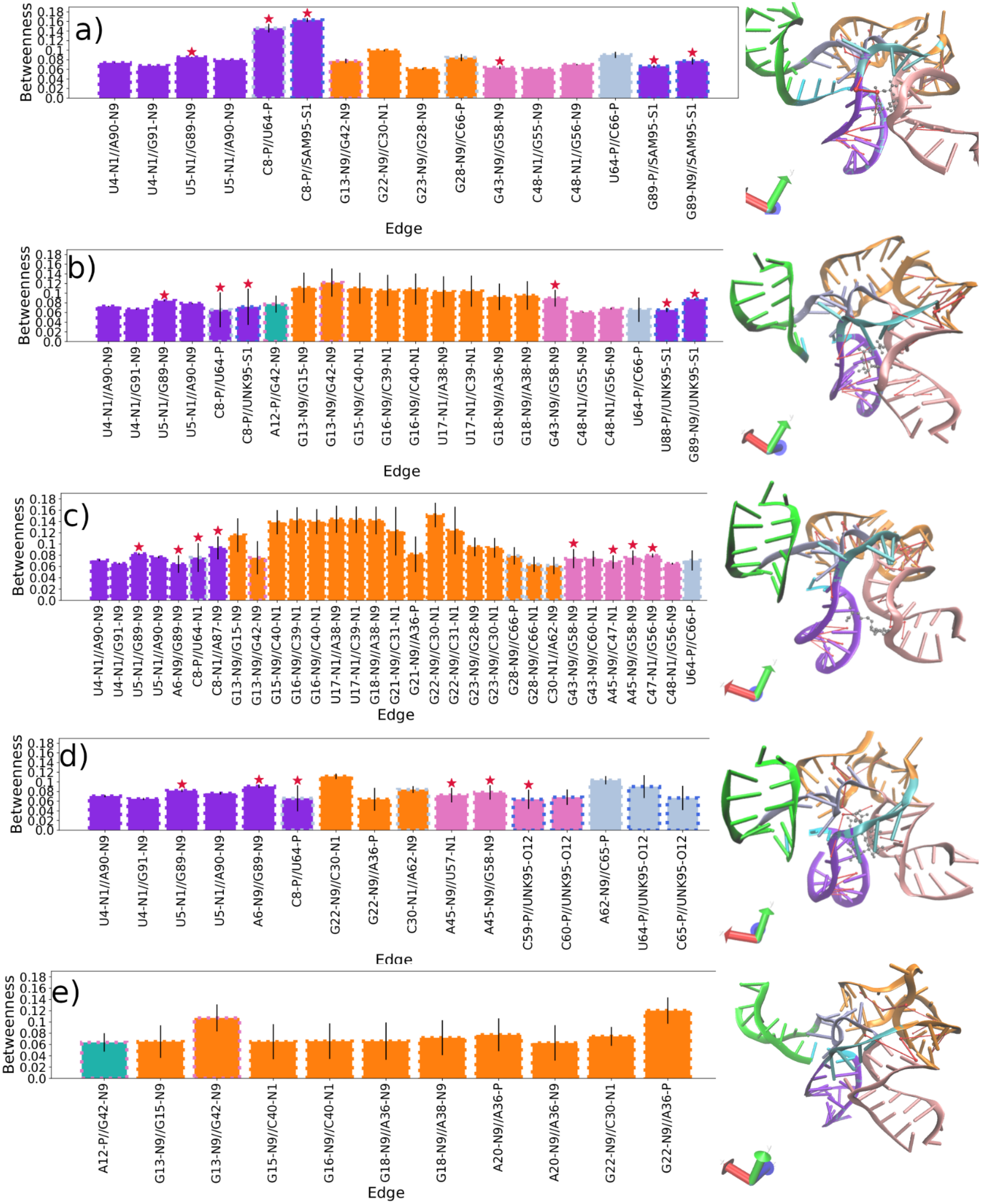
Average edge betweenness for node pairs having edges with betweenness >0.06. Bar plot face colors indicate the domain to which the first node is assigned, while dashed bar plot edge outline colors indicate the domain to which the second node is assigned. On the right hand we show the riboswitch complex, with domains colored according to the same color scheme, and the edges shown in the plot highlighted in red. Snapshots generated with VMD 1.9.4. Thicker edges have higher average betweenness. Error bars are calculated from the standard deviation across the five networks produced from the five different replicates of each system.

In **Fig. 11**, we visualize the eigenvector centrality of the nodes of the network that possess non-zero values. The eigenvector centrality, loosely, is a measure of the “influence” of a node over the entire network. In this context, a high eigenvector centrality indicates that the motions of that node exert a great deal of influence over the motions of other nodes in the network, controlling whether they are flexible or inflexible and how they move. From this, we observe that in the simulated native ligand-bound systems (SAMC, SAMD) and **only** in the native ligand-bound systems, all three nodes associated with the ligand are highly influential over the network (**Fig. 11a,b**, blue bars—values of about 0.3, 0.4), while **none** of the ligand nodes are highly influential in SAH or JS4. In addition, most of the other high centrality nodes are binding site nodes, including those associated with the P1 loop, the P3 loop, and the J12 region. Of particular interest are those nodes with the highest centrality in each region, which for SAMC and SAMD are the sidechain node of G11 (G11-N9) in J12, the sidechain node of G58 (G58-N9) in P3, and the backbone node G89 (G89-P) in the P1 loop. Conversely, in the SAH system and the JS4 system, most of the high-centrality nodes are associated with the P2 loop (**Fig. 11c,d**, orange bars). In particular, the highest-centrality P2 nodes in SAH are A20-N9 and G23-N9, while in JS4 it is A20-N9. The JS4 system shows less variability across replicates (**Fig. S9**, row 4) compared to the SAH system, which shows a pattern of shifting between influence that is localized at the tip of the P2 loop and influence located at the junction of the P2 and J34 regions (**Fig. S9**, row 5, **Fig. 11c**, ice blue bars, particularly A62-SC and C65-SC). In the SAH system, there is a small amount of influence exerted by the C8 residue in the binding site, as well as its neighboring node A9; this is negligible in the JS4 system. Finally, the unbound system demonstrates quite substantial differences in centrality locations across the different replicates (**Fig. S9,** bottom row, high error bars in **Fig. 11e**). Intriguingly, this corresponds to non-negligible but still lower influence appearing across domains that are important in SAM-bound systems (G11, **Fig. 11e**, turquoise, particularly G11-N9; P3 loop region, **Fig. 11e**, pink, with G58-N9 having a small but nonzero centrality) as well as domains that are important in SAH (A62 and C65, J34 region, **Fig. 11e**, ice blue) and in both SAH and JS4 (P4 loop, **Fig. 11e**, orange, particularly A20). On the other hand, little influence is ever localized in the P1 loop in the unbound state, though there is negligible but nonzero influence around A9. (For a more detailed discussion of the specific nodes implicated in the network, see Section **S**

**Fig. 11.**
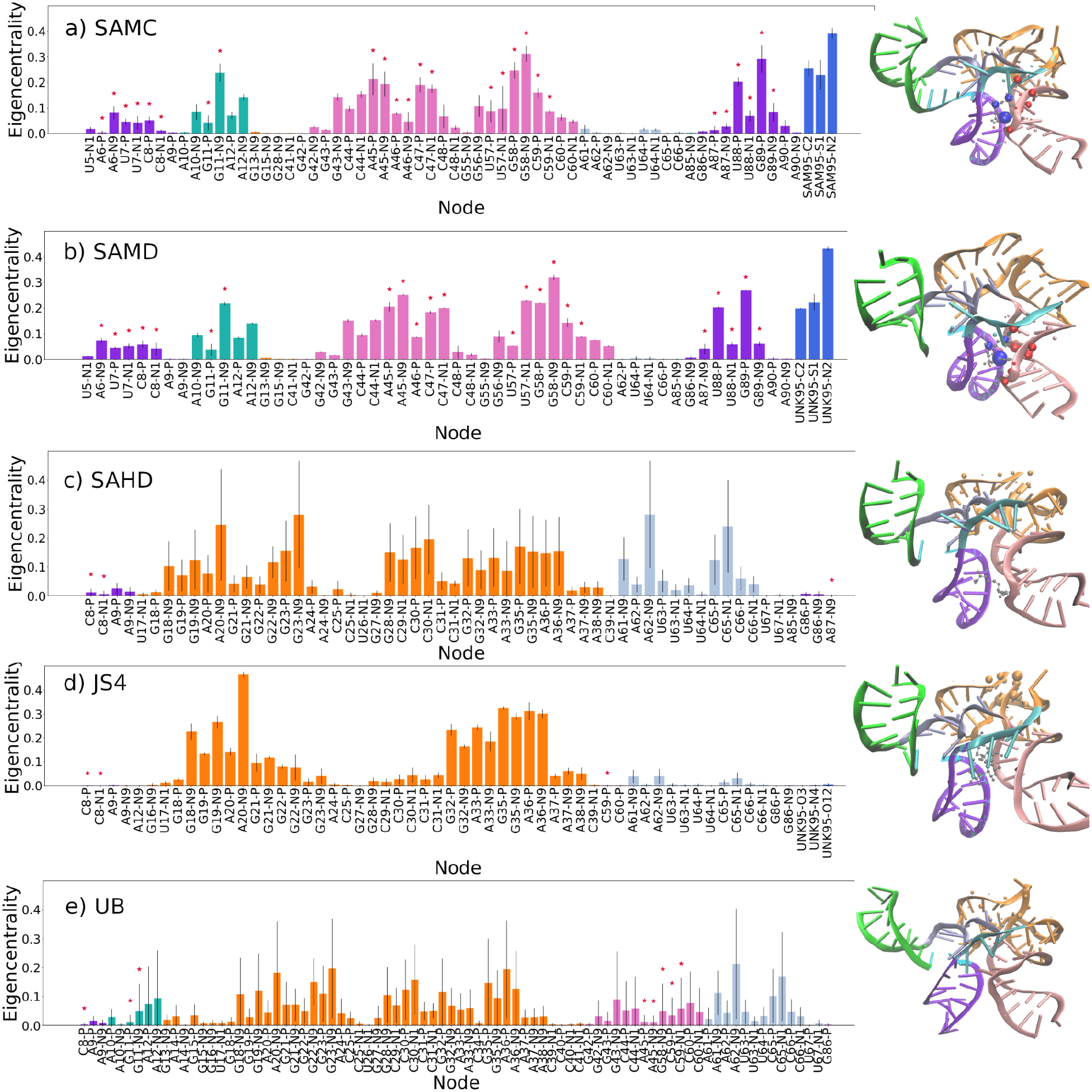
Average eigenvector centrality per nodes for nodes with >TOL values. Bar plot edge colors indicate the domain to which the node is assigned. Red stars indicate nodes that are part of the binding site. Snapshots on the right indicate the nodes of average higher betweenness as spheres with radii proportional to the eigenvector centrality. The complex is colored by the associated domain.

## 4. DISCUSSION

In this article, we used molecular simulations to model the dynamics of the interactions between the SAM riboswitch and different ligands. From the MD simulations, we extracted predictions about the local and global interactions that control riboswitch conformational behavior. Our local interaction predictions match observations from experimental static structures and our global interaction predictions provide a picture of the potential underlying dynamics. As expected, we observed strong ligand binding across all SAM-riboswitch replicates, but weaker binding/partial unbinding in the SAH system. As expected from docking predictions, we saw no reversal in the JS4 binding.

On a global level, we observed enhanced communication across the network for SAM-bound systems, as shown by larger communities, with one community of particular note in which two high-influence ligand nodes coordinate the P1 and P3 loop regions, including residues U7, U88, C47, and U57, and one community of particular note where the high-influence ligand tail node coordinates the J12 and P3 loop regions, including residues G11 and A45. The former community is notably split in all other systems, providing evidence of the SAM ligand directly enhancing P1 and P3 loop coordination, further bolstered by the high influence of the P3 loop over the network in SAM-bound systems. While we found evidence of global communication in JS4, the ligand was substantially less central and most network influence was concentrated in the P2 loop region, further from the P1 and P3 loops and without coordinating them. The SAH system, similarly, shows no strong influence of the ligand over the network, and a heightened concentration of network influence in the P2 and J34 regions. Most interestingly, the unbound system demonstrates that it has aspects of the SAM-bound system and of the SAH bound system, particularly evident in where eigencentrality appears in the network, though it shows little influence arising from the P1 loop.

Our predictions provide support for the following mechanistic rationale for experimental observations. In the native unbound state, two different modes are sampled, one in which the area near the P2 loop moves in a dynamically-coordinated fashion, one (less frequent) where the area near the P3 loop moves in a dynamically-coordinated fashion. When the native ligand is bound, it interacts strongly with the binding site residues, in particular with G11, U7, A87, U88, and C47. Hydrogen-bonding with the natively-inflexible G11 residue (inflexible in both the *unbound* and SAM-bound states, **Fig. S6**) functions as an anchor and to hold the ligand to the internal L1 loop in the J12 region, while coordination from charge interactions with U88 and pi-pi stacking with C47 locks the P1 and P3 loops together, and suppresses fluctuations in the P3 loop as well as the P1 loop. The ligand becomes a central focal point in a global network in the aptamer domain, which strongly stabilizes the terminator loop through stabilization of P1 by locking it to P3. SAH can stack with C47 but does not interact strongly with U88 or G11, thus failing to hold the P1 and P3 loops together and retaining more features of the unbound state, though it does partially stabilize P1 through its binding site interactions. Conversely, JS4 interacts strongly with U88, but contains no aromatic rings and does not interact with C47. In addition, the bulk of the molecule extrudes from the SAM binding site into the region nearer the P2 loop, leading to its stabilizing the alternate mode of the unbound state, in which the network of correlated dynamic motions is governed by the P2 loop, while the rest of the network moves in a less concerted and substantially more flexible fashion.

## 5. CONCLUSIONS

Overall, we have validated the hypothesis that SAM constrains the flexibility of the P1 loop compared to SAH. We have evidence that this may occur through locking of the P1 loop to the P3 loop via coordination between multiple different loop regions, with the most important direct interactions being those with residue G11 in the J12 region of the internal L1 loop, U88 in the P1 loop, and C47 in the P3 loop, which accords well with static experimental observations. Other important residues in this coordinated motion include A45 and G58 of the binding site in the P3 loop region. In addition, we have shown that, while the riboswitch can accommodate non-natural ligands in its binding pocket, for riboswitch modulation it is likely necessary to ensure interactions with the J12, P1, and P3 regions, though it is not clear whether the specific residues at hand are crucial. An instructive area for future research would be to attempt to design molecules that interact more strongly with nearby but differing residues, such as A12 in the J12 loop, C8 in the P1 loop, or A45 and G58 more directly. Our work adds to the body of literature cautioning against using docking results as a sole criterion for ligand discovery. Although we do not engage in this article with artificial intelligence techniques, the exquisite balance of specificity observed in ligand interactions with the riboswitch is also a more general caution against the use of less detailed computational techniques without complementary simulation or experiment, particularly when mechanisms of ligand action remain less-than-fully characterized.

## AUTHOR INFORMATION

### Author Contributions

The manuscript was written through the contributions of all authors. All authors have given approval to the final version of the manuscript. Specific contributions according to the CRediT taxonomy: Conceptualization - RAM, NM, and DE; Data curation - VN and RAM; Formal analysis - VN and RAM; Funding acquisition - RAM; Investigation - VN and RAM; Methodology - VN, RAM, and NM; Project administration - NM and RAM; Resources - NM and RAM; Software - NM and RAM; Supervision - NM and RAM; Validation - VN and RAM; Visualization - VN and RAM; Writing - original draft - VN and RAM; Writing - review & editing - all authors.

### Data Availability

Simulations and data analysis scripts are available on Zenodo at DOI https://doi.org/10.5281/zenodo.19135674. This includes structures (PDB files) and trajectories (in XTC format) of five different replicates for each system with water stripped out to make the file size manageable. We have also uploaded analysis scripts for the Rg, RMSD, RMSF, COM, h-bond analysis, pi-pi stacking, distances of sulfur atoms, and dynamical network correlation, as well as the raw data output of the dynamical network correlation analyses. Full trajectories are available from the authors upon reasonable request.

## ACKNOWLEDGMENT

The authors thank Prof. Srirupa Chakraborty, Prof. Claudine Gauthier, Prof. Nicolas Moitessier, and Dr. Janet Gaba for enlightening conversations and invaluable support.

