## Supplemental information - text and figures for "Molecular-level modulation of the SAM-I riboswitch drives global dynamical shifts"

### ASSOCIATED CONTENT

#### Supporting Information

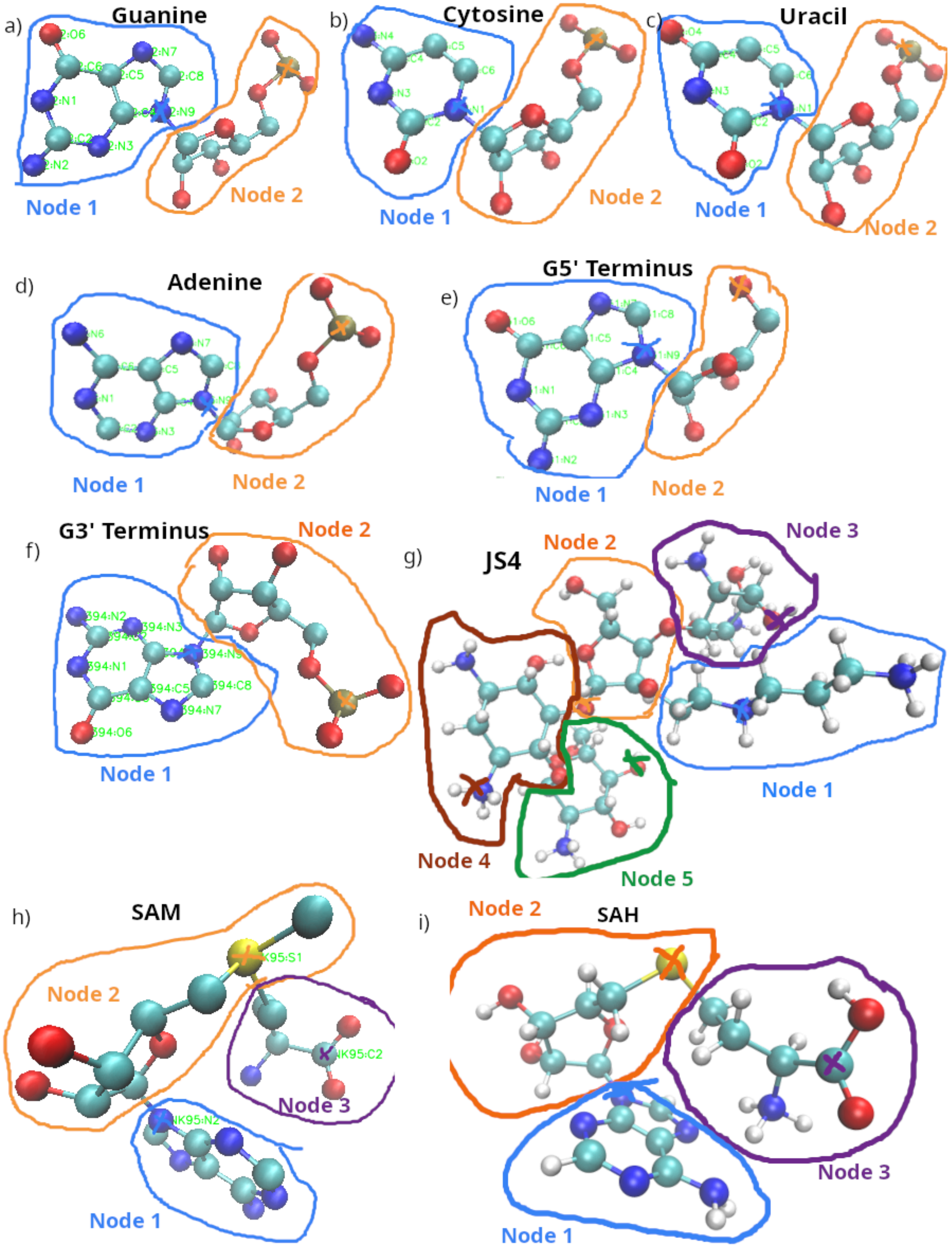
a)
Guanine
Node 1
Node 2
b)
Cytosine
Node 1
Node 2
c)
Uracil
Node 1
Node 2
d)
Adenine
Node 1
Node 2
e)
G5' Terminus
Node 1
Node 2
f)
G3' Terminus
Node 1
Node 2
g)
JS4
Node 1
Node 2
Node 3
Node 4
Node 5
h)
SAM
Node 1
Node 2
Node 3
i)
SAH
Node 1
Node 2
Node 3

**Fig. S1.** Node definitions for network analysis for (a) guanine, (b) cytosine, (c) uracil, (d) adenine, (e) G5’ terminus, (f) G3’ terminus (g) JS4, (h) SAM, and (i) SAH. Atoms belonging to one node are circled and the central atom is marked with an *X* of the same color.

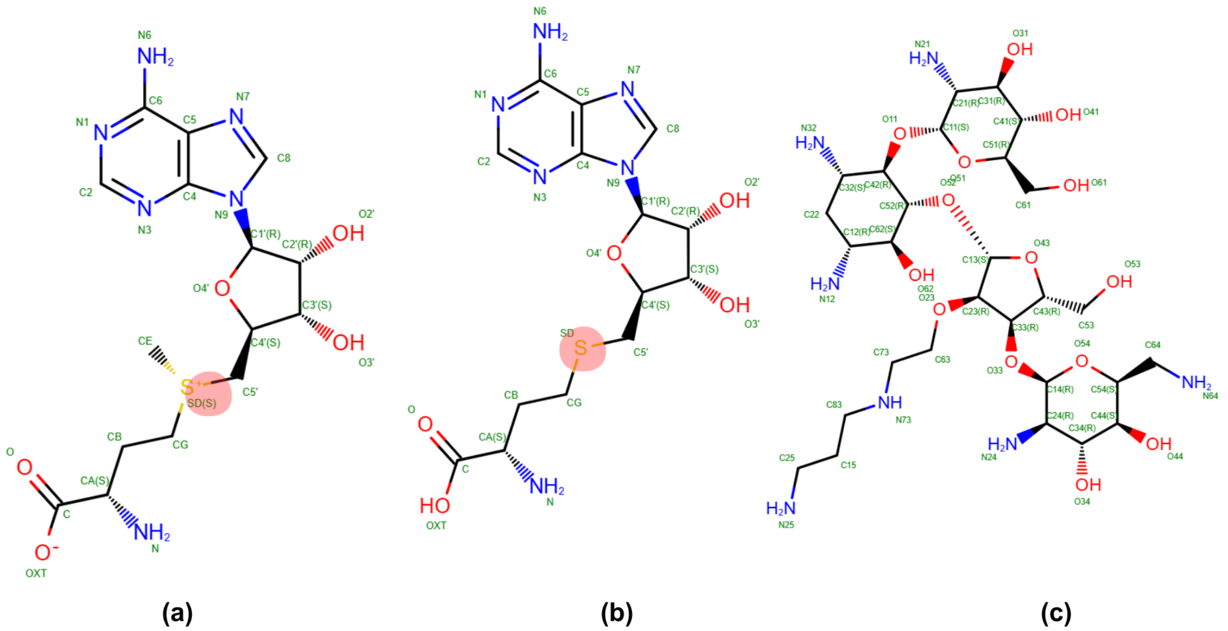


**Figure S2**: Chemical structures of studied ligands. a. SAM (positively charged sulfonium ion highlighted in red), b. SAH (highlighted sulfur neutral atom), c. JS4. Structures retrieved from the PDB database and visualized by PoseView[49]

### Section S1. Further Details of Virtual Screening

#### S1.1 Docking Procedure

For the virtual screening of small molecules, we first converted our 150 small molecule structures from SDF format to mol2 format using Conformational Optimization of Necessary Virtual Enantiomers, Rotamers, and Tautomers (CONVERT tool in FITTED). CONVERT generates an initial 3D structure by applying a local optimization algorithm to find a low-energy state. Using the automated PREPARE (Protein Rotamers Evaluation and Protonation based on Accurate Residue Energy, which also accepts RNA targets) tool in FITTED, we added missing hydrogens to the RNA target structure and defined the binding site of the target by selecting the co-crystallized SAM ligand in the 3GX5 structure. Given a co-crystallized ligand/target structure, FITTED uses a probe-based system to identify a grid of coordinates surrounding the relevant binding site. First, the software calculates the geometric center (centroid) of the selected reference ligand. Then, the PROCESS (Protein Conformational Ensemble System Setup) step uses a grid to generate beads of varying sizes that define the binding-site cavity. Unless otherwise stated, default values were used at all steps.

**Table S1:** SAM (self-docking) with RMSD and binding energy

| **Docked poses** | **RMSD (Å)** | **Energy value (kcal/mol)** |
|---|---|---|
| 1 | 1.18 | -98.919 |
| 2 | 1.22 | -109.650 (best pose) |
| 3 | 1.12 | -103.077 |
| 4 | 1.12 | -104.163 |
| 5 | 1.53 | -97.340 |

**Table S2:** SAH docked with binding energy

| Docked poses | Energy value (kcal/mol) |
|---|---|
| 1 | -99.339 |
| 2 | -98.036 |
| 3 | -108.138 (best pose) |
| 4 | -85.365 |
| 5 | -85.240 |

**Table S3:** JS4 docked with binding energy

| Docked poses | Energy value (kcal/mol) |
|---|---|
| 1 | -144.082 |
| 2 | -141.449 |
| 3 | -125.673 |
| 4 | -128.870 |
| 5 | -154.559 (best pose) |

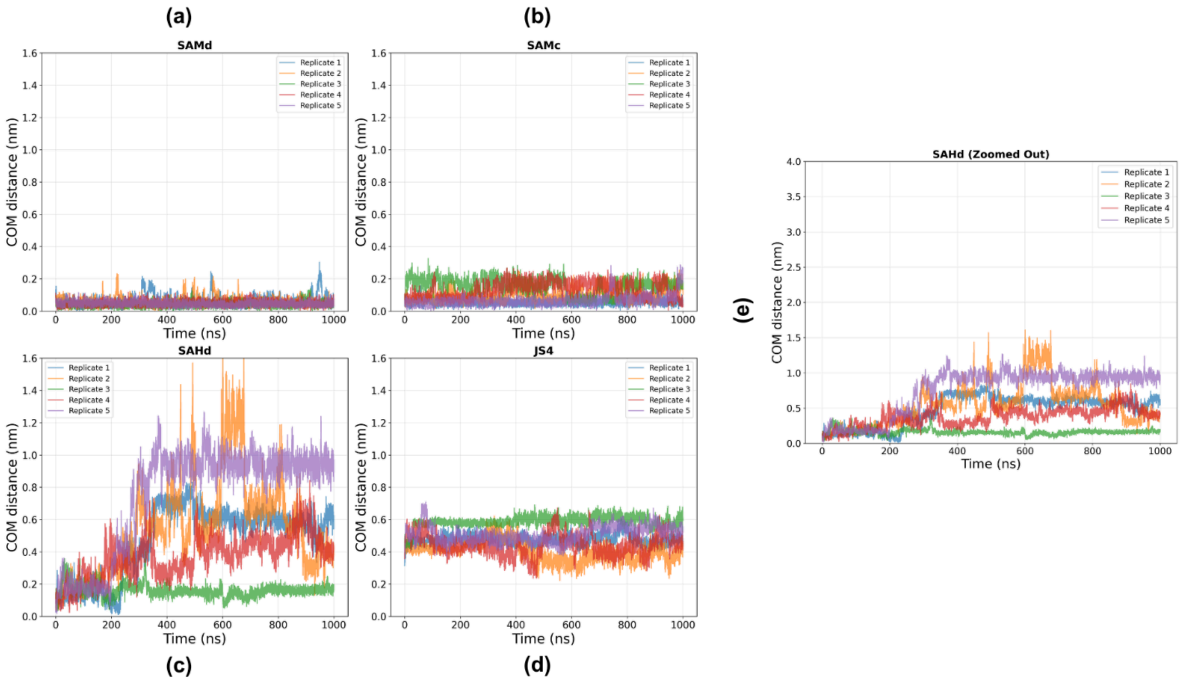


**Figure S2:** COM of all five replicas for each of the four RNA systems (a,b,c,d) and e. COM distance between ligand (SAH docked) and binding site vs time (zoomed out)

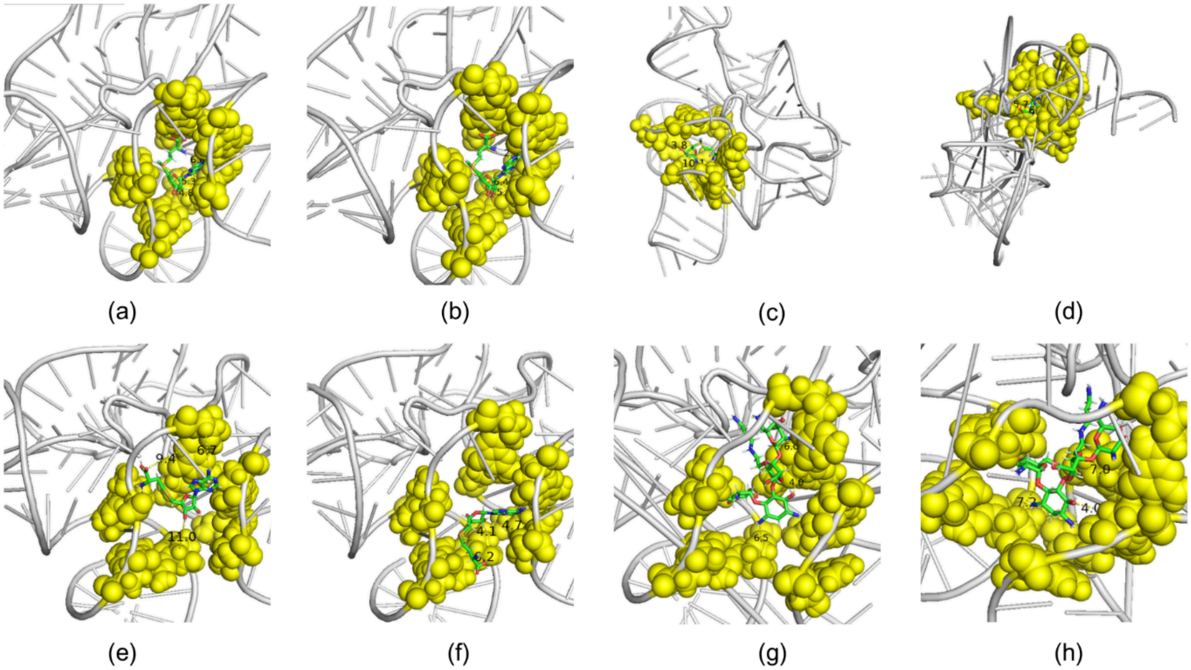

**Figure S3**: Representative MD simulation movie snapshots of four RNA-ligand systems. (a) RNA-SAMD at around 500 ns, (b) RNA-SAMD at around 900 ns, (c) RNA-SAMC at around 500 ns, (d) RNA-SAMC at around 900 ns, (e) RNA-SAHD at around 500 ns, (f) RNA-SAHD at around 900 ns, (g) RNA-JS4 at around 500 ns and (h) RNA-JS4 at around 900 ns. Snapshots rendered with Pymol.

#### Section S2 RNA-ligand complex is stable for all bound ligands

The radius of gyration ($R_g$) was calculated to assess the compactness of the RNA structure throughout the simulation. We calculated the mean and standard error across five independent replicates for each system **(Figure SXXa).** All systems start from an initially larger radius of gyration that drops rapidly, followed by clear evidence of equilibration within the first few nanoseconds. The SAMD system demonstrates a longer time for the decrease to occur (~50-100 ns), while the JS4 system equilibrates to a slightly more extended conformation (2.170 ± 0.009 nm versus ~2.13 nm for all other systems, cf. also Table S4).

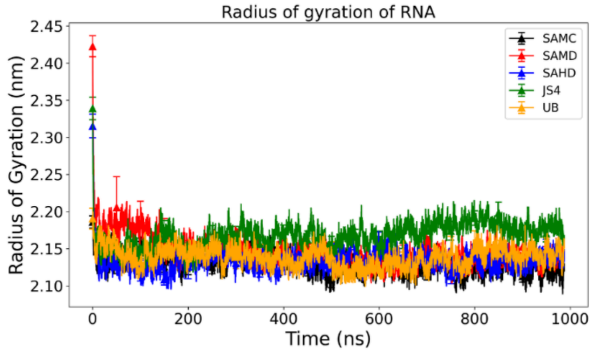


**(a)**

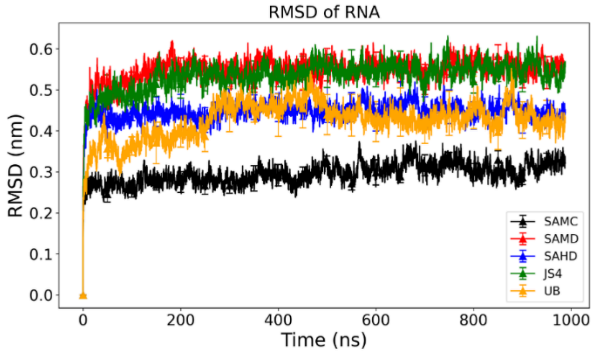


**(b)**

**Figure S4**: (a) Radius of Gyration of RNA and, (b) Root mean square deviation (RMSD) of

RNA for five systems (i) crystallized with the SAM ligand (SAMC), (ii) docked with the SAM ligand (SAMD), (iii) docked with the SAH ligand (SAHD), (iv) docked with the JS4 ligand (JS4), and (v) with the ligand removed (UB). Error bars represent the standard error taken across five independent replicates for Rg and RMSD.

**Table S4**: Radius of Gyration average ± SEM (five independent replicates, employing the final 900 ns to be certain of equilibration)

| **Systems** | **Average ± SEM** |
|---|---|
| SAM crystallized | 2.128 ± 0.003 nm |
| SAM docked | 2.143 ± 0.009 nm |
| SAH docked | 2.133 ± 0.009 nm |
| JS4 docked | 2.170 ± 0.009 nm |
| Unbound RNA | 2.138 ± 0.009 nm |

**Table S5**: RMSD average ± SEM (five independent replicates, employing the final 900 ns to be certain of equilibration)

| **Systems** | **Average ± SEM** |
|---|---|
| SAM crystallized | 0.299 ± 0.019 nm |
| SAM docked | 0.554 ± 0.026 nm |

| | |
|---|---|
| SAH docked | 0.450 ± 0.031 nm |
| JS4 docked | 0.542 ± 0.030 nm |
| Unbound RNA | 0.434 ± 0.047 nm |

RMSD calculations were performed to evaluate the structural deviation of the RNA from its initial conformation (**see Figure 3b**). The average RMSD with standard errors was calculated for RNA alone for all the complexes across five independent replicates. We observe that RNA in crystallized SAM shows the lowest RMSD (0.299 ± 0.019 nm over the last 900 ns), which indicates that RNA maintains a conformation closest to the reference structure, as we might expect, which provides *a posteriori* validation of our choice of force field and simulation parameters. The RMSDs of the unbound RNA and the RNA in the SAH (docked) complex are also stable and largely within error bars of the crystallized complex, with values of 0.434 ± 0.047 nm and 0.450 ± 0.031 nm, respectively. The RNA complexes docked with SAM and JS4, respectively, exhibit slightly higher RMSDs (0.554 ± 0.026 nm, 0.542 ± 0.030 nm after 900 ns, respectively). From the overall stability and comparative smallness of the RMSD values, however, we note there is unlikely to be substantial structural fluctuation apart from initial rearrangements.

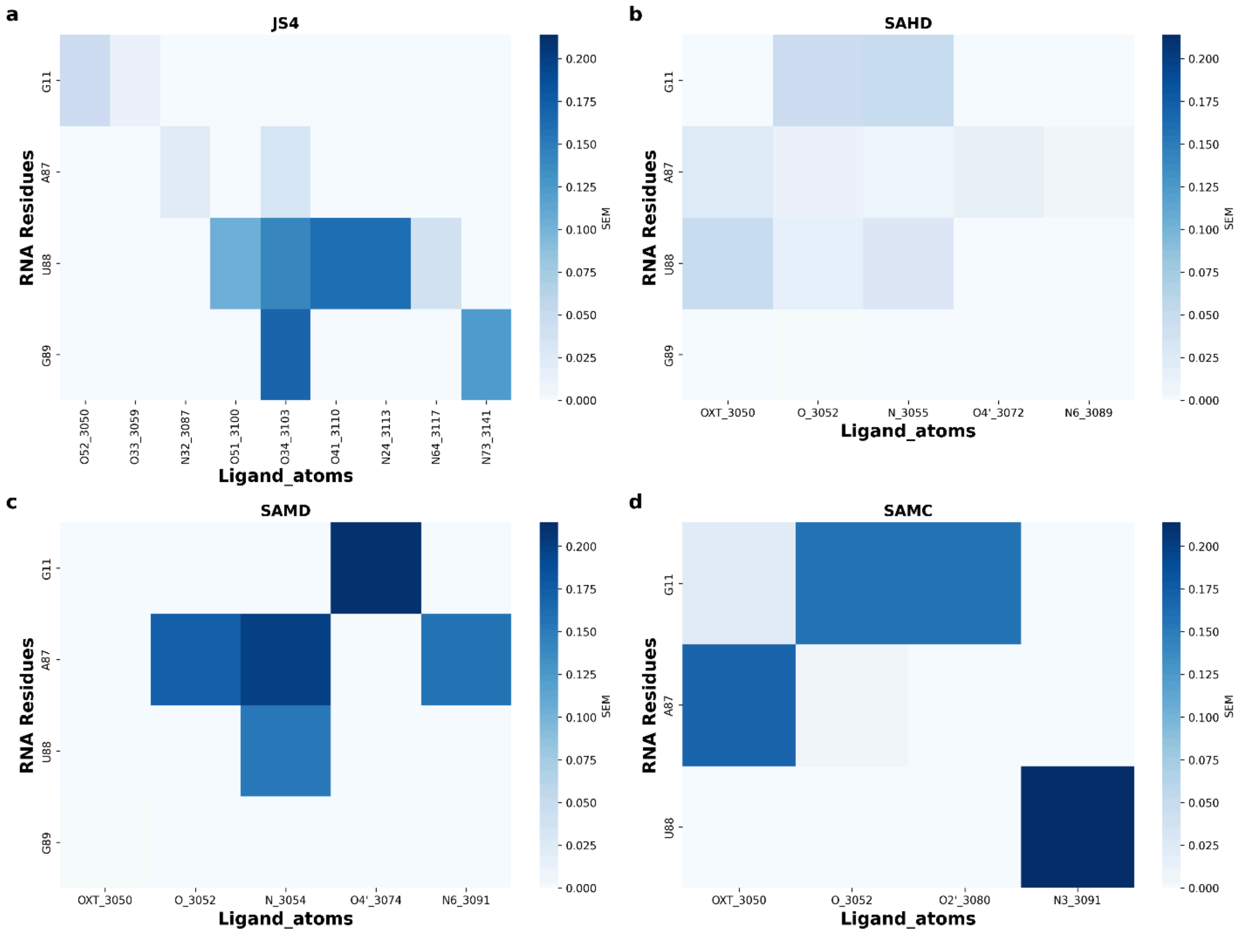


**Figure S5**: Hydrogen occupancy error heatmap for four RNA-ligand complexes. (a) Hydrogen bond occupancy error heatmap of RNA-JS4 complex (docked), (b) Hydrogen bond occupancy error heatmap of RNA-SAH complex (docked), (c) Hydrogen bond occupancy error heatmap of RNA-SAM complex (docked), (d) Hydrogen bond occupancy error heatmaps of RNA-SAM complex (crystallized)

**Table S6**: Average and standard error of the number of hydrogen bonds formed between RNA and ligands. Average taken over the final 900 ns.

| **Systems** | **Average ± SEM** |
|---|---|

| | |
|---|---|
| JS4 | 9.954361 ± 0.861996 |
| SAH docked | 3.438040 ± 0.434637 |
| SAM docked | 9.626175 ± 0.042317 |
| SAM crystallized | 8.164360 ± 0.523686 |

**Table S7.** Average occupancy (%) of pi-pi stacking interactions across five replicas between SAM and SAH ligands and the riboswitch binding site.

| Binding site residue | SAMC | SAMD | SAHD |
|---|---|---|---|
| A45 | 0.008 ± 0.009 | Not observed | 0.008 ± 0.003 |
| A6 | 0.008 ± 0.01 | Not observed | 1 ± 1 |
| A87 | Not observed | Not observed | Not observed |
| C47 | 99 ± 0.5 | 46 ± 1 | 10 ± 4 |
| G11 | Not observed | Not observed | 0.002 ± 0.002 |
| G58 | 0.8 ± 0.5 | 0.4 ± 0.1 | 0.4 ± 0.2 |
| G89 | 0.01 ± 0.007 | 0.005 ± 0.003 | 0.5 ± 0.3 |
| U57 | Not observed | 0.03 ± 0.01 | 0.10 ± 0.05 |
| U7 | Not observed | Not observed | 0.01 ± 0.01 |
| U88 | Not observed | Not observed | 0.001 ± 0.0006 |

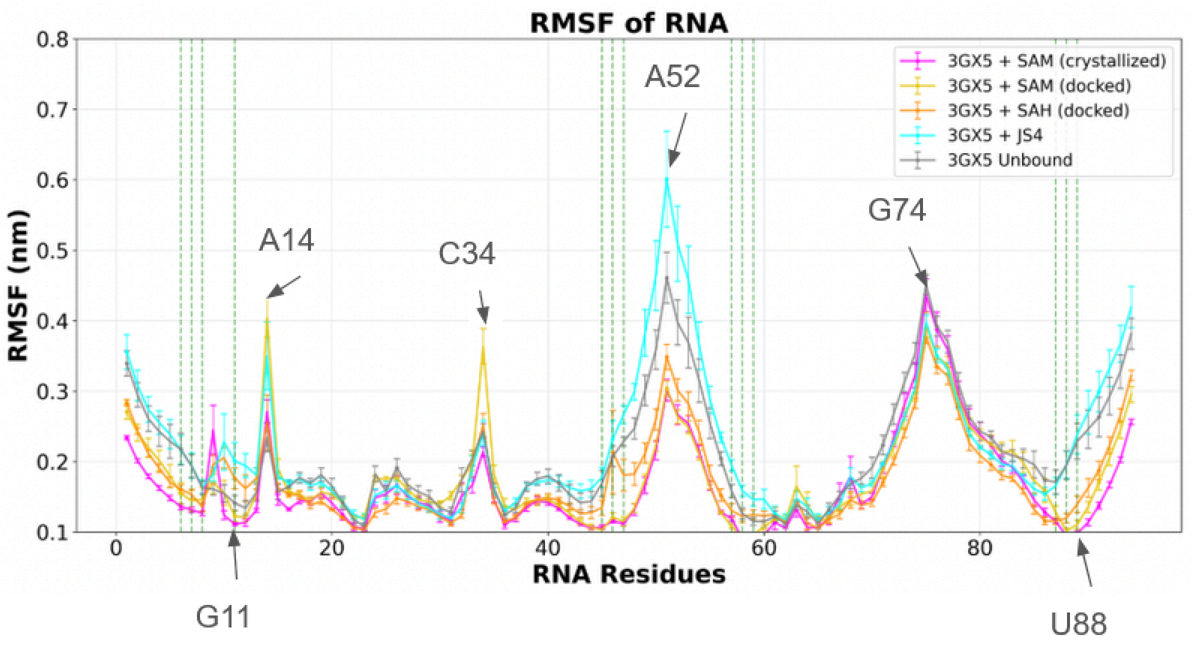


**Fig. S6.** Residue-wise RMSF of the riboswitch. Dashed green lines indicate binding site residues. Certain residues of interest are indicated with arrows.

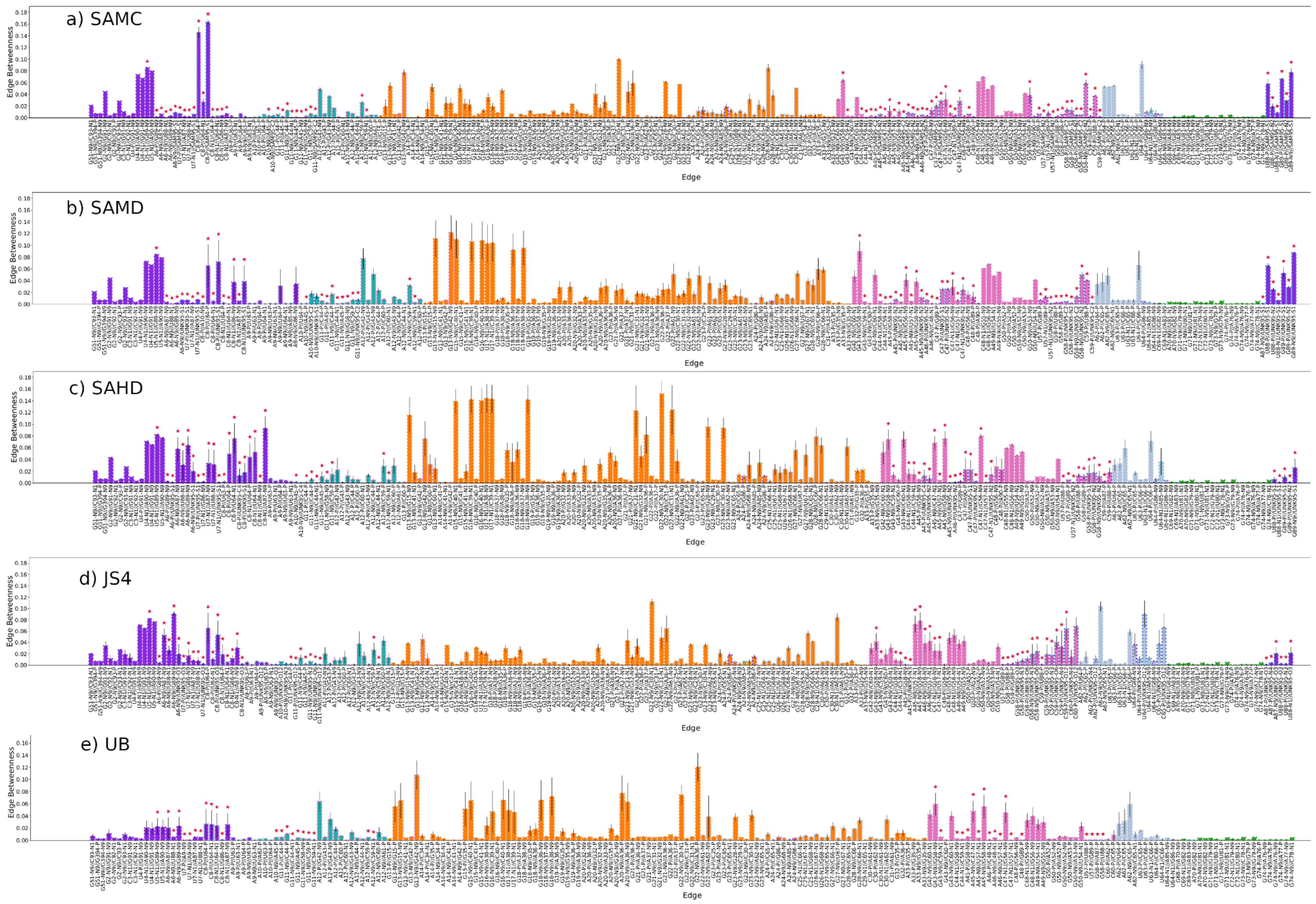

a) SAMC
b) SAMD
c) SAHD
d) JS4
e) UB
Edge Betweenness
Edge

**Fig. S7.** Average edge betweenness for all node pairs with non-zero betweenness. Bar plot face colors indicate the domain to which the first node is assigned, while dashed bar plot edge outline colors indicate the domain to which the second node is assigned. Error bars are calculated from the standard deviation across the five networks produced from the five different replicates of each system.

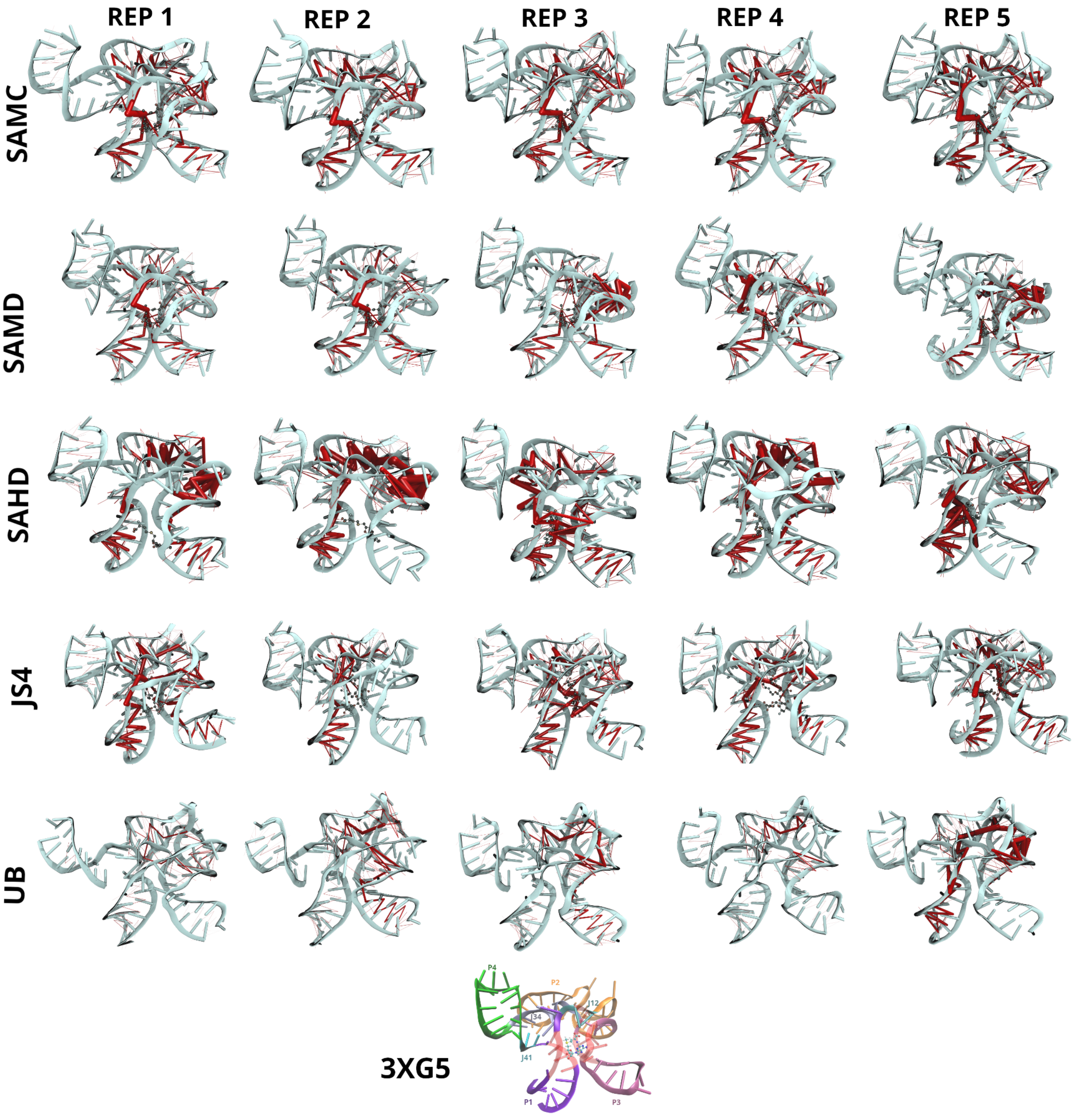

**Fig. S8.** Visualization of edge betweenness for the network constructed from each replicate using the Dynetan python package and VMD. Edges with non-zero betweenness are indicated as red cylinders, and edge weight is indicated by cylinder thickness.

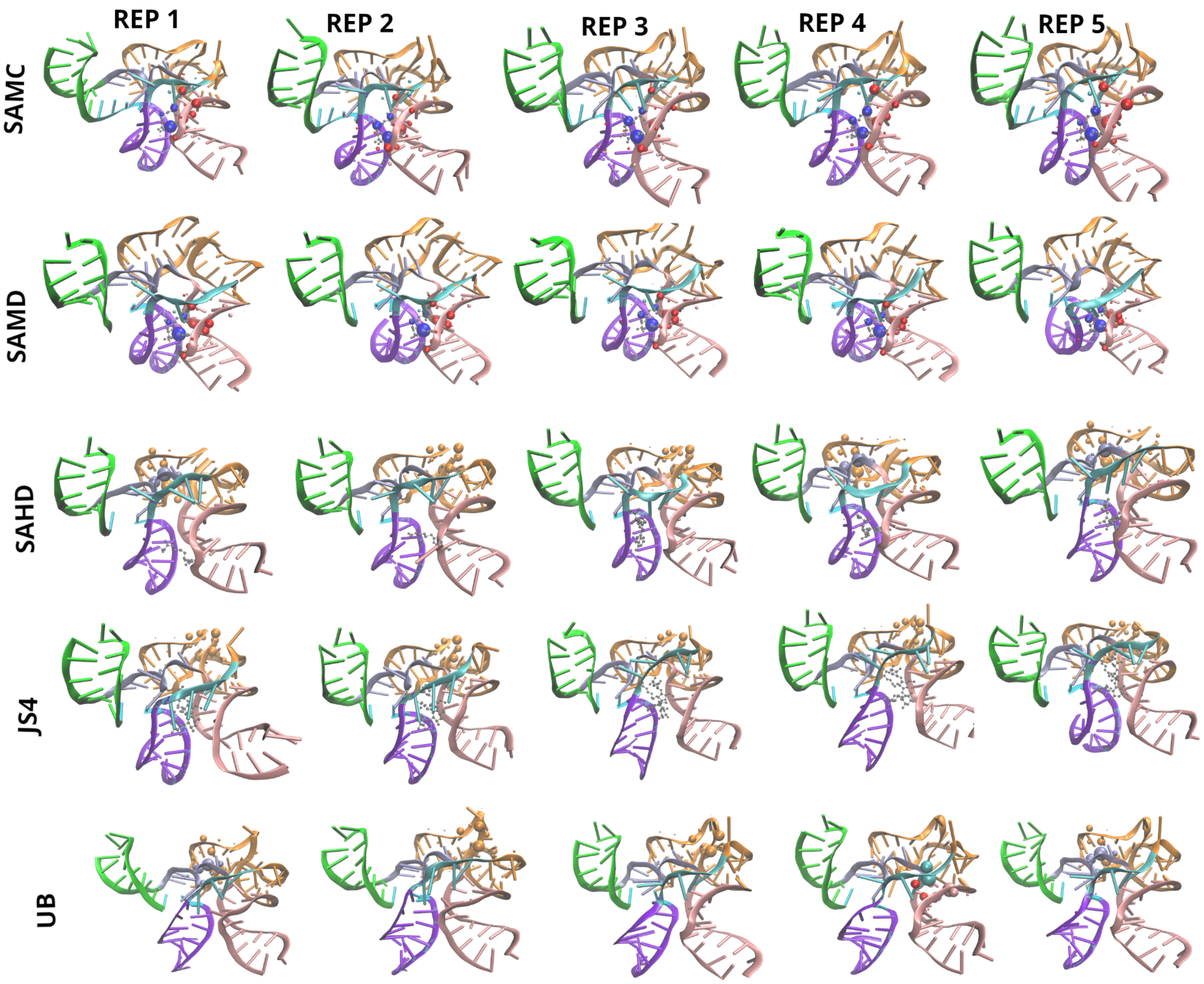


**Fig. S9.** Visualization of node eigenvector centrality for the network constructed from each replicate using VMD. Spheres indicate nodes with non-negligible eigenvector centrality, and the radius of the sphere is proportional to the weight. Colors indicate domains. Blue nodes are ligand-associated nodes, and red nodes are associated with the experimentally-determined binding site.

**Section S3. Participation of individual residues in local and global interactions.** Drawing all this together, a complex, dynamical picture of binding site interactions emerges, validating and complementing experimental evidence. SAM h-bonds with residue G11 in the J12 loop via both the methionine tail and the ribose ring, while SAH only interacts via its methionine tail. This is similar to experimental observations highlighting G11, but in those (static) observations, its proximity to the tail is the only one highlighted. G11, additionally, demonstrates high influence over the SAM network in all cases, with a small amount of influence over the unbound network. Residue U88, which is part of the P1 loop and is substantially more flexible in the unbound state than when bound to SAM or SAH, maintains as suggested a steady distance to the charged sulfonium in SAM but not to the neutral sulfur in SAH; however, it also forms h-bonds with the methionine tail and to a lesser extent the pyrimidine ring of SAM, and the methionine tail of SAH, also to a lesser extent. It also h-bonds with JS4. Despite strong hydrogen bonding with the ligand in the JS4 system, it does not appear as influential in the network in any way. U7 also maintains a steady distance to the charged sulfonium in SAM but not the neutral sulfur in SAH. Despite U7 being slightly closer to the sulfonium than U88, its substantially lower eigencentrality suggests that of the two, the interactions with U88 are the more important to the network overall, perhaps due to its additional h-bonding. On the other hand, U7 is directly adjacent to C8, through which dynamical information flows to U64 in the J34 loop in SAMC, SAMD, and SAHD, but which has high informational flow to the *ligand* only in the SAM-bound systems, though it is not central to the network. Residue C47, which is in the P3 loop, indeed does display persistent pi-stacking with SAM, and also to a small extent with SAH. In the SAM-bound systems alone, it is found in a community with U7/U88, and only in this situation exerts moderately high influence over the dynamical network formed by the complex, suggesting importance in modulating the P1/P3 loop coordination. Other residues that emerge from simulation are A87 and G89, both in close proximity to U88 and generally belonging to the same community. A87 forms h-bonds with the SAM methionine tail and pyrimidine ring, while G89 forms h-bonds with JS4. While A87 is not implicated further as a particularly important residue in the network, G89 emerges as a very high-influence node in the SAM-bound systems, but not in the JS4 system.
